# Orally available designed miniproteins inhibit enterotoxigenic *Bacteroides fragilis* pathology by blocking toxin receptor binding

**DOI:** 10.64898/2026.06.22.733822

**Authors:** Pooja Srinivas, Victor Adebomi, Susan M. Markiewicz, Kang Wang, Denise Chac, Kristofer Lindenauer, Nishoni Huber, Zhihang Tao, Phat Luong, Stephen A. Rettie, Maisie W. Smith, Asim K. Bera, Alex Kang, Hannah Nguyen, Maika Schneider, Yaxi Wang, S. Brook Peterson, Min Dong, Ana A. Weil, Gaurav Bhardwaj, Joseph D. Mougous

**Author notes:** To whom correspondence should be addressed: Email –. These authors contributed equally (Pooja Srinivas, Victor Adebomi).

## Abstract

Toxigenic bacterial infections in the gut are a significant contributor to the global burden of disease. Advanced tools for protein and live biotherapeutic engineering offer potentially transformative strategies for treating such diseases, while avoiding the collateral effects of traditional antibacterials. Here we used de novo protein design to identify inhibitors of the metzincin family protease *Bacteroides fragilis* toxin (BFT). These inhibitors, which bind distal to the active site, interfere with toxin-mediated E-cadherin cleavage and downstream proinflammatory signaling by blocking claudin-4 receptor binding. We tested the inhibitors as disulfide-stabilized variants administered directly to the cecum or in drinking water, as well as through in situ secretion by an engineered live biotherapeutic. Across these delivery modalities, the inhibitors successfully neutralized the toxin and effectively prevented BFT-associated gut pathology, including tumor formation. These results highlight the potential of de novo designed proteins as precise, non-antibiotic interventions to mitigate bacterial toxin-driven disease in the gut.

## Introduction

Bacterial toxins are often major drivers of disease, promoting tissue damage, immune evasion and dysregulation, and transmission. Therefore, toxin neutralization represents a promising alternative to traditional antibiotics for treating bacterial infections. Small molecule-based approaches targeting toxins have largely focused on suppressing production, with few examples of compounds that directly inhibit toxin activity^1–3^. In contrast, there are numerous examples of biologic approaches directly targeting toxins, predominantly utilizing monoclonal antibody (mAb) and nanobody platforms^4^. One notable success in this arena is bezlotoxumab, a human IgG1 mAb targeting *Clostridioides difficile* toxin B, which is FDA approved for preventing recurrent *C. difficile* infection^5^. This drug and others in pre-clinical development bode well for toxin neutralization strategies targeting pathogens resident in the gastrointestinal tract^6–8^. Such therapies hold the promise of mitigating disease while preserving microbiome composition and reducing the selective pressure for antibiotic resistance associated with conventional antibiotics.

Despite the promise of protein-based toxin neutralization strategies for gastrointestinal pathogens, many challenges remain. Oral drug delivery to the colon is complicated by chemical and enzymatic degradation in the gastrointestinal tract, limiting drug concentrations at sites of infection and necessitating alternative delivery approaches^9^. For example, bezlotoxumab targets a toxin produced in the colonic lumen yet requires intravenous administration^5^. Though not unique to therapeutic biologics in the GI tract, development and manufacturing costs are also an impediment. In cases where toxin neutralization does not accompany pathogen eradication, costs could be compounded by the need for long-term administration. Indeed, despite its efficacy, bezlotoxumab was ultimately withdrawn from the market^10^.

Enterotoxigenic *Bacteroides fragilis* (ETBF) is a clinically important intestinal pathogen that requires the secreted metalloprotease *Bacteroides fragilis* toxin (BFT) to promote disease^11,12^. BFT is a metzincin-family toxin encoded within a pathogenicity islet absent from non-toxigenic *B. fragilis*^13^. Depending on the population studied, approximately 5-30% of individuals are colonized with ETBF^14^, and it is associated with diarrheal disease and inflammatory bowel disease (IBD)^15,16^. ETBF is more prevalent in children with diarrhea compared to healthy controls, and one study found it in 51% of adult patients with ulcerative colitis (n=35), compared to 1.5% of matched controls (n=60)^14,16,17^. ETBF-associated inflammation is also implicated in colon cancer; multiple studies report higher ETBF incidence or *bft* gene presence (3-12X) in colon cancer patients relative to control cohorts^18–20^.

The molecular mechanisms by which BFT promotes ETBF pathogenesis are incompletely understood. The toxin is comprised of a type II signal peptide, inhibitory prodomain, and catalytic domain^11^. Following secretion, the prodomain is removed by the cell surface-associated cysteine protease fragipain, releasing the active toxin^21^. Mature BFT cleaves the E-cadherin ectodomain, disrupting adherens junctions and compromising barrier integrity in colonic epithelia^22^. This enhances epithelial permeability and promotes cytoplasmic accumulation of β-catenin^23,24^, predisposing individuals that harbor or acquire adenomatous polyposis coli (APC) mutations to colonic neoplasia^22,25^. These pro-oncogenic effects of BFT are amplified through NF-κB activation, inducing inflammatory mediators including interleukin-8, which can act as an autocrine growth factor^26–30^.

Recent advances in artificial intelligence (AI)-based *de novo* protein design have enabled the generation of mini-protein binders (“minibinders”). These compact proteins, typically fewer than 100 residues, can achieve high-affinity target recognition while maintaining thermal and proteolytic stability^31,32^. Unlike conventional biologics, minibinders can be produced in their active form in highly scalable and relatively inexpensive heterologous hosts, such as *E. coli*^31,33^. This also theoretically permits their direct integration into engineered live biotherapeutic products (LBPs). In the gut environment, LBPs offer the capacity for long-term therapeutic delivery at the sites of infection^34,35^. In this study, we developed BFT minibinders that act distal to the active site at the prodomain interface to inhibit toxin receptor binding at nanomolar concentrations. These inhibitors ameliorate toxin-induced pathology when supplied directly to the cecum or when consumed in drinking water. We further developed LBP strains that secrete the minibinders and demonstrate their efficacy in treating ETBF catalyzed tumorigenesis. This work identifies de novo designed proteins with therapeutic potential in individuals harboring ETBF, and it establishes a generalizable experimental framework for toxin neutralization in the gut.

## Results

### De novo design identifies prodomain interface binding inhibitors of BFT

*B. fragilis* strains can carry one of three BFT isoforms, which share ∼90% sequence identity and are each associated with disease. We targeted our designs against BFT-1, the most prevalent subtype in human ETBF isolates (∼65%)^36^. Prior to proteolytic activation, BFT is autoinhibited through a prodomain aspartate switch that extends into the catalytic site and coordinates the active-site zinc ion (Figure 1A) ^37^. As an initial minibinder design strategy, we sought to recapitulate this inhibitory mechanism. Our minibinder design pipeline consisted of three-steps (Figure 1B): (1) backbone generation using RFdiffusion with conditioned denoising trajectories (lengths spanning 45-65 amino acids, n=10,000), (2) amino acid sequence design using ProteinMPNN (two sequences per backbone) and (3) candidate filtering via AlphaFold2 (AF2) and Rosetta-based metrics (see Methods)^32,38–41^. After applying our filtering criteria, we retained 140 designs, which we further diversified by partial denoising diffusion combined with LigandMPNN^39,42^. Further filtering yielded a final library of 14,730 candidates for experimental screening.

**Figure 1.**
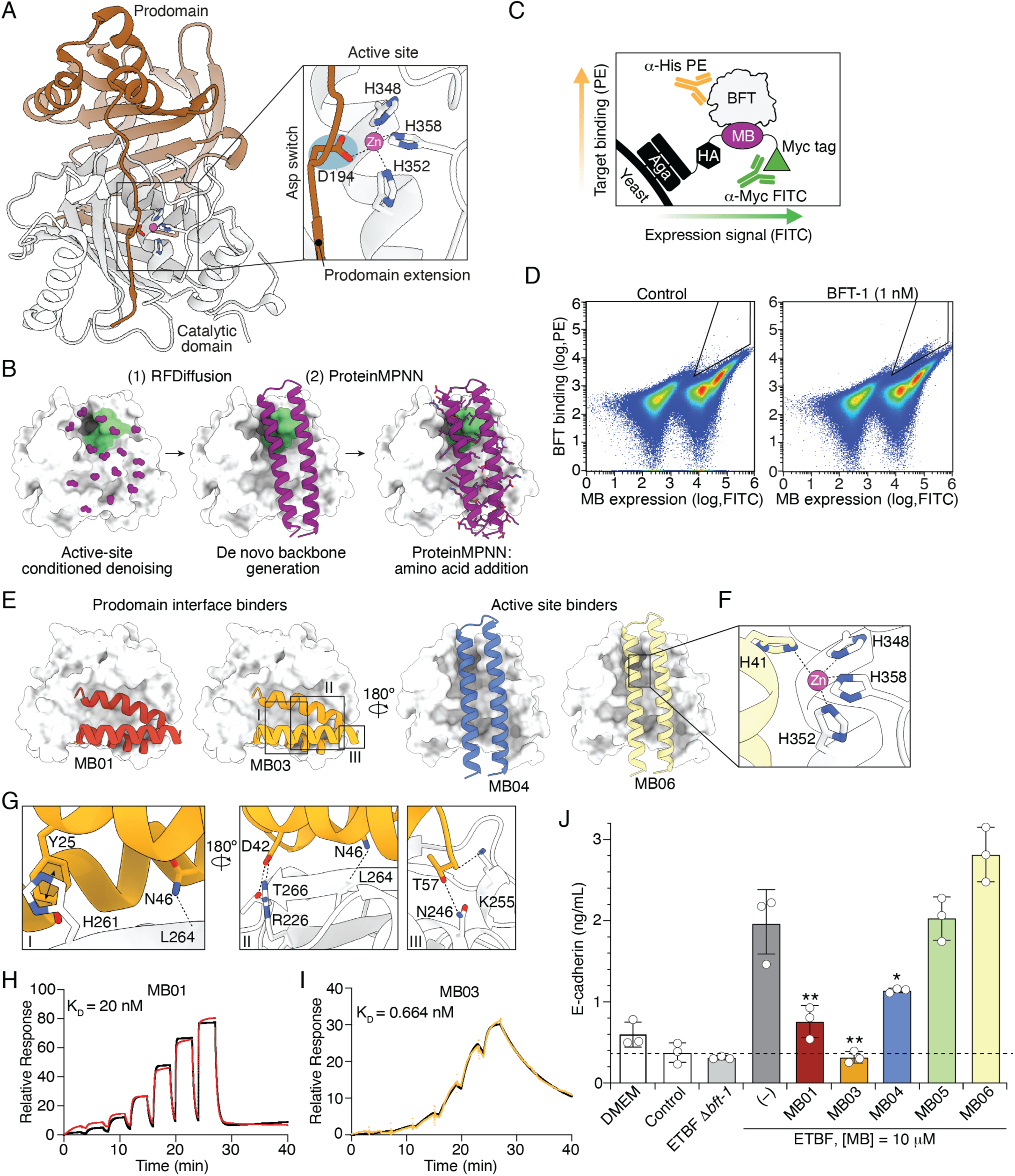
Identification of prodomain interface binding inhibitors of BFT. A) Ribbon diagram depiction of the BFT-1 structure (8H3Y), with inset depicting aspartate switch-mediated inhibition of catalytic residues by the prodomain extension. B) Minibinder design strategy, including backbone generation via conditioned denoising with RFDiffusion^32^ (1) and amino acid sequence introduction using ProteinMPNN^39^ (2). The BFT-1 catalytic domain is depicted in surface mode (white). C) Schematic showing FACS-based yeast surface display strategy for BFT-targeting minibinder enrichment. A yeast library displaying C-terminal Myc-tagged minibinders is incubated with BFT-1–H_8_, then stained with fluorescently labeled antibodies targeting Myc and 8×His to indicate MB expression and BFT-binding, respectively. FITC, fluorescein isothiocyanate; PE, phycoerythrin; MB, minibinder. D) Representative pseudocolor density plots of FITC and PE fluorescence from the yeast display minibinder library, following two rounds of FACS-based enrichment for BFT-1 binding and exposure to 1 nM BFT-1 (right) or the control treatment. Antibody staining was applied as depicted in (C), and black lines indicate gating applied during sorting. E-G) AF3-based predictions of the interaction sites identified for representative enriched minibinders, with BFT-1 residues in the active site (F) or prodomain interface (G) predicted to make contacts with minibinders shown in insets. H, I) SPR analysis of the binding affinity of the indicated minibinders for immobilized BFT-1–H_8_. K_D_ values indicate binding affinity calculated from this analysis. Colored traces indicate measured data; black curves indicate corresponding fitted data. J) Soluble E-cadherin detected in the cell supernatant from HT-29 cells exposed to ETBF SCS and the indicated treatments. Data represent means and standard errors and are representative of multiple experiments conducted. Asterisks indicate values significantly different from ETBF spent cell supernatant treatment (1-way ANOVA with Dunnett’s multiple comparisons test, *p<0.001, **p<0.0001).

To identify BFT binders, we displayed our library of designed miniproteins on yeast and subjected cells to enrichment in two rounds of fluorescence-activated cell sorting (FACS) (Figure 1C, and Supplemental Figure 1A,B)^31^. Following enrichment, a BFT-1 titration series (1 nM-1 μM) was used to identify high-affinity candidates (Figure 1D, Supplemental Figure 1C). Structural modeling of the enriched binders predicted two distinct binding modes (Figure 1E, Supplemental Figure 2). As expected, one class targeted the active site (Figure 1F). Interestingly, a smaller subset of binders – derived from a single non-catalytic site binding candidate identified in our initial design – was predicted to engage the prodomain interface^37^. These prodomain interface binders (PIBs) are predicted to form extensive interactions with BFT-1 (Figure 1G). Given their favorable computed binding metrics and high enrichment in yeast surface display screening (see Methods), these prodomain-interface designs were advanced alongside catalytic-site binders.

We selected the most enriched BFT minibinder candidates, three helix bundle PIBs (MB01-MB03), from the 1 nM BFT titration sort for further characterization (Supplemental Figure 1D). The catalytic site binders were significantly less enriched, so we selected the top three, each comprised of two α-helices, for further testing (MB04-MB06, Supplemental Figure 2). MB02 exhibited low solubility that precluded analysis. Surface plasmon resonance (SPR) assays revealed no detectable BFT binding for MB04-MB06. In contrast, MB01 and MB03, which share 79% sequence identity, bound BFT-1 with high affinity (K_D_=20 nM and 0.66 nM, respectively, Figure 1H,I, Supplemental Table 1), consistent with the favorable computed binding metrics of these prodomain interface binders. These affinities are similar to those of other de novo designed minibinders for their targets and significantly exceed those of previously reported nanobodies occluding the BFT active site^33,37,43–45^.

We next assessed the inhibitory capacity of the minibinders by quantifying E-cadherin release from the immortalized colonic epithelial cell line HT-29, a commonly used assay for measuring BFT activity^22,46^. We used filtered spent cell supernatant from ETBF or ETBF Δ*bft-1* (as a control) as a source of BFT in these experiments (Supplemental Figure 3A). At a concentration of 10 μM, both PIBs significantly inhibited E-cadherin release by BFT – approximating the cleaved E-cadherin levels in control samples not exposed to the toxin (Figure 1J, Supplemental Figure 3B). Despite our inability to detect BFT binding by SPR, we found MB04 also inhibits E-cadherin release by the toxin. We speculate that in the case of this minibinder, our SPR measurements were confounded by sensor chip binding. Neither MB05, MB06, nor a binder designed against an unrelated protein target, inhibited E-cadherin release, consistent with their absence of detectable binding to BFT (Figure 1J, Supplemental Figure 3C).

### PIBs exhibit high affinity for BFT and potently inhibit its activity

The performance of PIBs motivated us to generate second-generation minibinder libraries targeted to this surface of BFT. We designed and screened two additional libraries: one restricted to fewer than 66 residues and another spanning larger sequence lengths (66-88 residues) (Supplemental Figure 4A-F). Yeast surface display sort enrichment values, estimated binding affinities determined from BFT titration sorts, fold characteristics, and overall geometry were used to select candidates for experimental analysis (see Methods) (Supplemental Figure 4G, Supplemental Figure 5). Three candidates we selected, MB07, MB15 and MB22, are strongly predicted PIBs and exhibit high affinity for BFT-1 (K_D_ = 1.45 nM to 11.5 nM) (Figure 2A-D, Supplemental Figures 6A-E, Supplemental Table 1). We also assessed the binding of these minibinders – along with the top first generation PIB MB03 – to BFT using a precipitation approach (Figure 2E). This orthogonal method provided additional evidence for their capacity to tightly associate with the toxin. Based on these results, we proceeded with further mechanistic and functional assays using these four PIBs.

**Figure 2.**
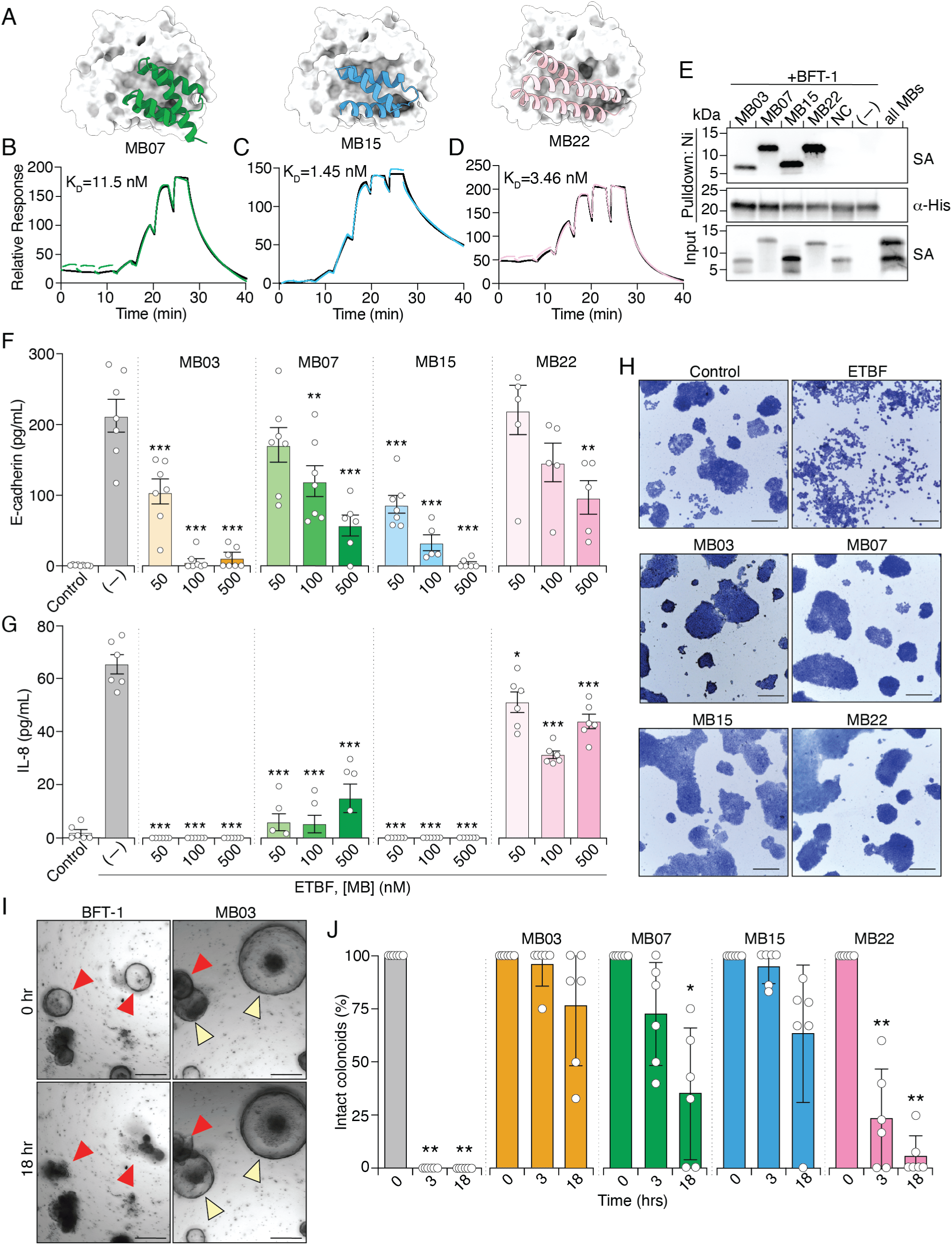
PIBs exhibit high affinity for BFT-1 and robustly inhibit BFT activity. A-D) Structural models (A) and SPR-based BFT-1 binding affinity assays (B-D) for second generation PIB minibinders. Colored traces indicate measured data; black curves indicate corresponding fitted data. E) Western blot analysis of biotinylated minibinders precipitated with BFT-1–H_8_. SA, streptavidin; NC, negative control binder designed against a different target. F-G) Released E-cadherin (F) or IL-8 (G) detected in the cell supernatant from HT-29 cells exposed to BFT-1-containing SCS and varying concentrations of the indicated minibinders or a vehicle control. H) Giemsa-stained HT-29 cells exposed to ETBF-1 SCS and the indicated minibinders or a vehicle control. Scale bar = 200 μm I, J) Murine colonoid integrity following treatment with purified BFT-1 alone (100 nM) or with indicated minibinders (10 mM). Red arrows indicate ruptured colonoids, yellow arrows indicate intact colonoids. Data in F, G and J represent means and standard errors and are representative of multiple experiments conducted. Scale bar = 250 μm. Asterisks indicate values significantly different from the ETBF SCS treatment (F,G: 1-way ANOVA with Dunnett’s multiple comparisons test; J: paired t-test with Bonferroni correction compared to 0-hr baseline; *p<0.01, **p<0.001, ***<0.0001).

To determine the relative potency of the four PIBs, we tested their ability to inhibit BFT-mediated E-cadherin cleavage from HT-29 cells across a range of concentrations. MB03 and MB15 inhibited the toxin at the lowest concentration tested (50 nM), whereas MB07 and MB22, required higher concentrations, 100 nM and 500 nM, respectively, to achieve significant inhibition (Figure 2F, Supplemental 6F-H). Reports show that E-cadherin cleavage by BFT promotes NF-κB activation and the release of IL-8, driving inflammation in the gut^29,47,48^. It also catalyzes cytoskeletal reorganization, leading to cell rounding and detachment^49^. Thus, as secondary measures of minibinder efficacy, we measured their capacity to inhibit IL-8 secretion and monolayer disruption catalyzed by BFT. Remarkably, MB03 and MB15 reduced IL-8 below the detection limit of our assay at 50 nM, suggesting their potency could exceed that inferred solely by E-cadherin measurement (Figure 2G). MB07 and MB22 also inhibited IL-8 secretion at 50 nM; however, likely due to trace contaminants (e.g. endotoxin), measurements at higher concentration were confounded by weak IL-8 induction from the protein preparations themselves. At 1 μM, each minibinder strongly or completely inhibited HT-29 aggregate dissolution and cell rounding caused by BFT (Figure 1H, Supplemental Figure 6I). Finally, to better model native gut epithelia, we assessed the efficacy of PIBs in a colonic organoid model derived from C57BL/6J mice (Figure 2I). Treatment with purified BFT-1 (100 nM) in the absence of minibinder induced rupture of all colonoids within three hours (Figure 2J, Supplemental Figure 7). In contrast, intact colonoids remained following BFT treatment in the presence of each PIB (10 μM) beyond the 18 hours. MB03, MB07 and MB15 exhibited particularly robust protection. Mirroring our other phenotypic assays, MB22 provided protection against the toxin, but below that of the other PIBs.

### PIBs inhibit BFT catalysis and receptor binding

The distance of the surface of BFT predicted to interface with PIBs from the active site of the toxin suggests that this class of minibinders achieve their potency through a mechanism distinct from competitive inhibition. Though improbable given the high structural prediction confidence metrics, such as AF TM-scores (0.88-0.93 average across the four PIBs) (Supplemental Figures 2,5), we first considered whether the predicted interface of the PIBs could be incorrect. To test this, we introduced amino acid substitutions in MB03 at selected predicted BFT-interacting residues (Figure 3A). Pulldown assays and bio-layer interferometry analyses with these mutants yielded results anticipated by AF modeling (Figure 2B, Supplemental Figure 8A). The introduction of charged residues at central sites in the predicted interface strongly diminished binding (I32D, M36K, V49D), whereas more conservative substitutions weakened binding (Y25A, N46A, V50A) and substitutions at distinct sites acted additively (Y25A, N46A) (Figure 2B, Supplemental Figure 8A). Although we cannot rule out that the same residues contribute to binding on another surface of the toxin, these data support the AF-predicted structure of PIB–BFT complexes.

**Figure 3.**
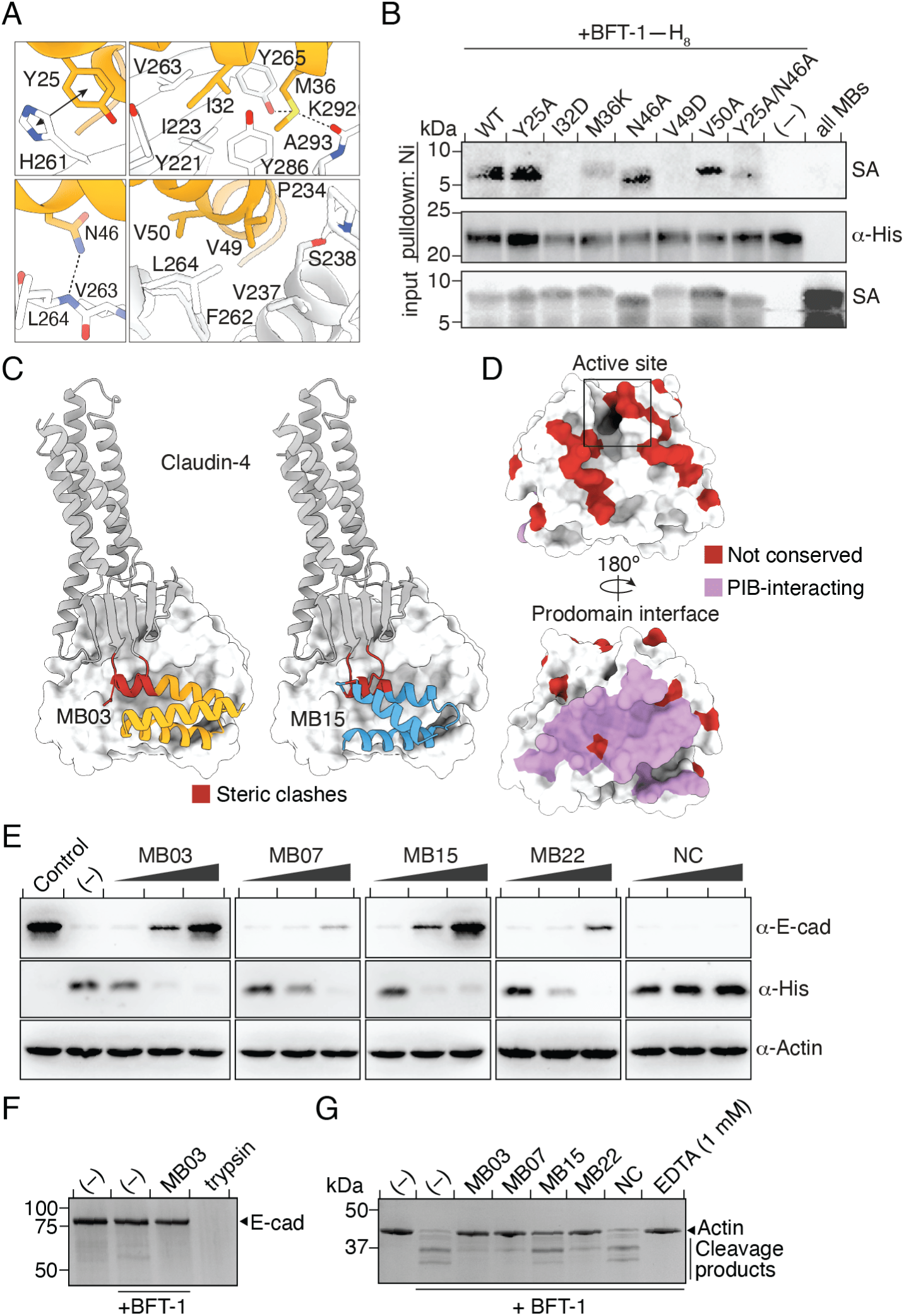
PIBs inhibit BFT interaction with the receptor claudin-4 and reduce BFT catalytic activity. A) Structural model zoom-in showing predicted interacting residues from MB03 (orange) and BFT-1 (white). Arrow, aromatic interaction; dashed lines, electrostatic; not indicated, hydrophobic. B) Western blot analysis of biotinylated MB03 with the indicated substitutions precipitated with BFT-1–H_8_. C) Model of the predicted interaction between claudin-4 (grey) and BFT (white) with the predicted binding site of two PIBs indicated. Those PIB residues predicted to clash with claudin-4 interactions are highlighted. D) BFT-1 structure (8H3Y) indicating residues not conserved in BFT-2 and their location relative to the PIB interaction interface. E) Western blot analysis of proteins extracted from HT-29 cells alone (control) or incubated with BFT-2 (5 nM) and increasing concentrations of the indicated PIBs (1:1, 10:1 or 100:1 PIB:BFT-2 molar ratio). NC, negative control minibinder designed against a different target. F, G) Coomassie-stained SDS-PAGE analysis of products generated from five hours incubation of purified BFT-1 (20 nM) with 2 μM of E-cadherin (F) or actin (G) and minibinders (1 μM) added as noted. NC, negative control as in E.

We recently identified the tight junction protein claudin-4 as a receptor for BFT^50^. Knockout of this polytopic membrane protein in HT-29 cells abrogates BFT cell binding and BFT-mediated cleavage of E-cadherin. A structure of the BFT–claudin-4 complex has not been determined and AF modeling suggests claudin-4 may bind a surface of BFT that partially overlaps with that of our PIBs; therefore, we hypothesized that PIBs might inhibit BFT by interfering with receptor binding (Figure 3C, Supplemental Figure 8B,C). To test this, we incubated BFT with wild-type and claudin-4 knockout cells in the presence and absence of PIBs. Notably, we performed these experiments with BFT-2. This isoform occurs at lower prevalence than BFT-1, but exhibits higher activity *in vitro*, causes more severe disease in animal models, and is more strongly associated with colorectal cancer^19,36,51,52^. Of the 25 amino acid differences between BFT-1 and BFT-2, only four are located outside of the catalytic face of the toxin and the two of these within the prodomain interface are conservative substitutions that are not predicted to impact PIB binding (Figure 3D, Supplemental Figure 8D). Indeed, we found that when present at 50 nM, each PIB reduced claudin-4 dependent BFT-2 cell binding to nearly background levels (Figure 3E, Supplemental Figure 8E). Western blot analysis of E-cadherin on the same cells indicated that higher minibinder concentrations are required to achieve an equivalent degree of inhibition of its cleavage by the toxin, suggesting that the small fraction of BFT that binds claudin-4 becomes less accessible to the minibinders (Figure 3E). Together, these results provide strong evidence that PIBs prevent BFT-mediated cleavage of E-cadherin from colonic epithelial cells by blocking the interaction between BFT and claudin-4.

Although PIB inhibition of receptor binding could fully account for the BFT inhibitory activity of the minibinders, this does not preclude that the minibinders also inhibit the enzymatic activity of the toxin. Studies typically report a failure to robustly detect *in vitro* cleavage of E-cadherin by BFT^22,53,54^. Despite optimizing BFT enzymatic conditions using the model substrate actin, we similarly detected only faint apparent E-cadherin cleavage products *in vitro* (Figure 3F, Supplemental Figure 9A,B)^55,56^. We also noted that a recent study reporting direct cleavage of E-cadherin by BFT incubated >50 μM toxin at near stoichiometric levels with the substrate in order to achieve significant turnover^43^. Therefore, it was necessary to use the non-native substrate actin to more conclusively assess the impact of PIBs on BFT catalysis^56^. Our results show that 1 μM of each PIB significantly attenuates BFT-mediated actin cleavage (Figure 3G, Supplemental Figure 9C). MB15 inhibited the toxin to a lesser extent than the other PIBs, possibly reflecting the higher K_off_ of this minibinder (Supplemental Table 1). Taken together, these data demonstrate that PIBs block BFT binding to claudin-4. They also suggest that PIBs further interfere with BFT activity by allosterically inhibiting catalysis. Since receptor binding appears to be required for BFT-mediated E-cadherin cleavage, it is unclear what degree direct inhibition of catalysis plays in the overall efficacy of PIBs. Nevertheless, these dually operating mechanisms of inhibition may slow the emergence of resistance.

### Development of tools for *in vivo* administration of PIBs

The therapeutic potential of PIBs relies upon identifying a practical and effective route of delivery. Though there is evidence in mice that inhibiting BFT via nanobodies can reduce pathology associated with IBD, this required intravenous administration of 200 mg/kg four times weekly^43^. Such a regimen is likely prohibitive for treatment of a chronic GI infection like that of ETBF. Oral availability of a BFT biotherapeutic offers numerous potential advantages, including lower cost, improved efficacy by targeting to the site of infection, and compatibility with long-term treatment; however, even for therapeutics that are ultimately encapsulated, this delivery route requires overcoming the acidic and proteolytic environments of the upper gastrointestinal tract^9^.

We reasoned that LBPs engineered to secrete the PIBs could represent one solution to overcoming the challenges of oral administration (Figure 4A)^57,58^. We evaluated *Bacteroides ovatus* ATCC 8483, a candidate next-generation probiotic with high secretory capacity, and non-toxigenic *B. fragilis* NCTC 9343, which occupies the same niche as ETBF, as candidate chassis (Supplemental Figure 10A)^59–62^. Both strains are native constituents of the human microbiome with high colonization fitness and genetic tractability^62^. In an effort to optimize PIB production, we screened combinations of two *Bacteroides* promoters (P_BT1311_, P_BfP1E6_) and ribosome binding sites (rpiL*, RBS8) that were previously shown to confer high levels of heterologous protein expression^62,63^. To facilitate secretion, we selected a type II signal peptide derived from *B. thetaiotaomicron* identified in a comprehensive study by Sirk and colleagues to support exceptionally robust export of protein cargo from *Bacteroides* (BT_0294)^62^. The resulting constructs were stably integrated in single copy into the attB2 site on the chromosomes of *B. fragilis* and *B. ovatus*^64^. Western blot analyses revealed substantially higher levels of MB03 secretion from *B. ovatus* than *B. fragilis*, with MB03 detectable across all constructs and maximal levels achieved using the P_BfP1E6_-RBS8 combination (Figure 4B).

**Figure 4.**
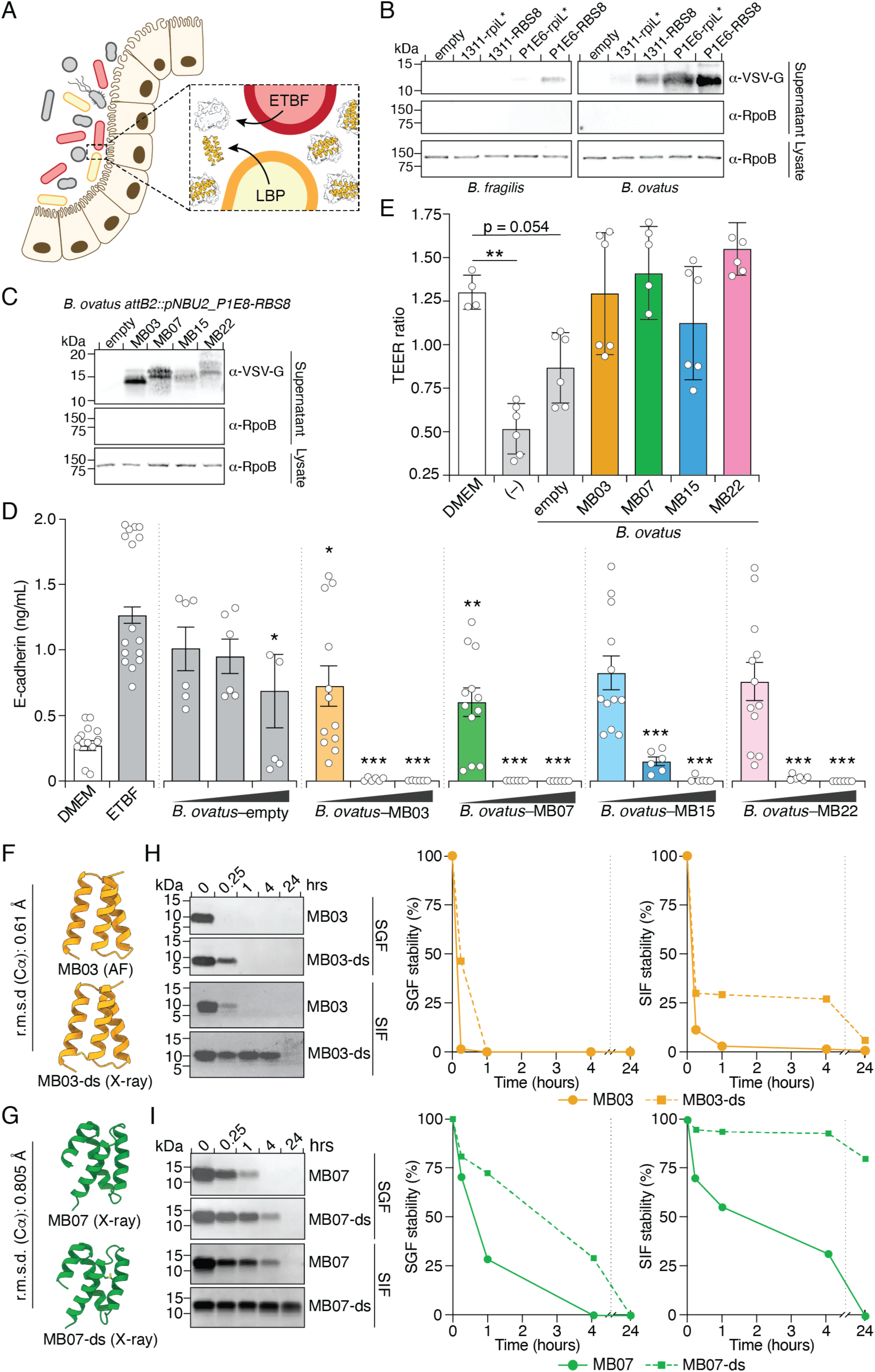
Development of tools for oral delivery of PIBs. A) Schematic depicting the approach of using PIB-secreting LBPs to inhibit BFT activity *in vivo*. B, C) Western blot analysis culture supernatant or cell lysate from the indicated *Bacteroides* strains engineered to secrete VSV-G–MB03 (B) or the indicated VSV-G–tagged PIBs (C) under the control of the indicated promoter and ribosome binding site combinations. RpoB staining was used a cytoplasmic protein control. D) Released E-cadherin detected in the cell supernatant from HT-29 cells exposed to media alone or ETBF and an increasing concentration of the indicated PIB-secreting strain (1:10, 1:1 or 10:1 initial ratio of ETBF:*B. ovatus*). Data represent means and standard errors from four biological replicates performed in triplicate. Asterisks indicate E-cadherin levels significantly less than in ETBF alone treated cells. E) Final over initial TEER ratio obtained from confluent HT-29-MTX-12 cells grown in transwells and incubated five hours with media alone, or ETBF (OD_600_=1.5) and the indicated PIB-secreting strains of *B. ovatus* (OD_600_=2.5) supplied to the apical chamber^66^. Data represent means and standard errors from six replicates. F,G) AF model or experimentally determined structure of the indicated minibinders and H,I) (left) SDS-PAGE analysis of the indicated PIBs incubated for the indicated intervals in simulated gastric or intestinal fluid (SGF, SIF), with the relative abundance of the full-length PIBs quantified by densitometry in ImageJ (right). Asterisks indicate values significantly different from the ETBF treatment (D) or DMEM (E) (1-way ANOVA with Dunnett’s multiple comparisons test, *p<0.5, **p<0.01, ***p<0.0001).

Next, we used E-cadherin release from HT-29 cells co-incubated with ETBF and our engineered LBP strains to evaluate the functional potential of secreted PIBs. *B. ovatus* secreting MB03, but not a strain carrying a control construct, abolished ETBF-mediated E-cadherin cleavage (Supplemental Figure 10B,C). In line with our secretion results, *B. fragilis* bearing MB03 did not significantly diminish E-cadherin cleavage under the conditions of this assay.

Similar bacterial burdens across conditions indicate that the absence of protection is driven by insufficient minibinder secretion rather than bacterial abundance (Supplemental Figure 10C). Given these results, we proceeded to generate *B. ovatus* LBPs expressing each of the PIBs under the control of P_BfP1E6_-RBS8 and fused to the signal peptide of BT_0294 (Figure 4C). Strains secreting MB03 and MB07 showed the greatest potency in co-culture assays with HT-29 cells, significantly inhibiting BFT when present at one-tenth the level of ETBF (Figure 4D, Supplemental Figure 10D-F). However, all LBPs conferred marked reduction in cleaved E-cadherin levels when present at equivalent levels as ETBF.

The most severe clinical manifestations of ETBF infection, including anaerobic sepsis, are driven by BFT-mediated disruption of the adherens junctions^21^. This disruption leads to cell rounding and an increase in paracellular permeability, allowing ETBF and other opportunistic gut bacteria to translocate from the lumen into the underlying lamina propria and bloodstream^65^. We further assessed the capabilities of the LBPs by measuring transepithelial electrical resistance (TEER) across HT-29-E12-MTX monolayers^66,67^. As expected, ETBF induced a marked decrease in TEER in a manner dependent on BFT (Figure 4E, Supplemental Figure 11). Co-culture with control *B. ovatus* did not prevent this loss of barrier integrity whereas all minibinder-secreting strains preserved TEER at levels comparable to untreated controls (Figure 4E, Supplemental Figure 11C). Together, these data demonstrate that PIBs secreted by *B. ovatus* can strongly suppress BFT-dependent cellular phenotypes associated with ETBF colonization and thereby support the notion that LBPs represent a viable strategy for oral administration of this class of BFT inhibitors.

A second way we envisioned achieving oral availability of the PIBs was to enhance their stability in the GI tract. The introduction of disulfide bonds into the minibinder scaffold has proven an effective means of promoting their passage through the acidic and proteolytic environment of the upper GI tract^33^. Disulfide bonds were not included the initial PIB design; therefore, we used Rosetta-based tools to identify candidate positions and evaluate energetic and functional parameters associated with incorporating disulfide bonds in candidate PIBs. We selected, expressed and purified the top disulfide-constrained version of each PIB (MB03,07,15,22-ds), all of which retained high affinity binding to BFT and the capacity to inhibit BFT-mediated E-cadherin cleavage from HT-29 cells (Figure 4F,G, Supplemental Figure 12A-C). X-ray crystal structures of MB03-ds and MB07-ds confirmed their high similarity to the predicted or experimentally determined structures of MB03 (0.61 Å r.m.s.d.) and MB07 (0.805 Å r.m.s.d), respectively (Figure 4F, G, Supplemental Table 2). Finally, we examined the stability of these disulfide-containing variants using simulated gastric and intestinal fluids (SGF, SIF), which mimic both the pH and proteolytic environment of their respective compartments (Figure 4H,I, Supplemental Figure 12D)^68^. With the exception of MB15-ds, each disulfide variant exhibited substantially improved stability in these solutions relative to their native PIB (Figure 4H,I, Supplemental Figure 12D). This was particularly pronounced for MB07-ds and MB22-ds; over a 24-hour incubation in SIF these variants exhibited no discernable degradation and each was readily detectable after a four-hour incubation in SGF. The stability of MB03-ds, MB07-ds and MB22-ds in SGF and SIF approximates that of other orally available protein therapeutics, suggesting that direct ingestion of these proteins or encapsulated versions thereof could represent a viable route of inhibiting BFT in the gut^33^.

### PIBs prevent BFT-linked pathology in a murine model

With tools in hand for oral PIB administration, we sought to determine whether PIB delivery to the gastrointestinal tract can influence BFT-mediated pathology in a murine model. As a first test for the feasibility of this approach, we employed a model in which purified BFT is directly injected into the mouse cecum^50^. Four hours following BFT injection, histological staining revealed extensive epithelia cell shedding and edema, consistent with previous reports. Strikingly, co-injection of BFT with a 10-fold excess of MB03-ds or MB15-ds, the two most inhibitory minibinders, abolished BFT-linked tissue damage (Figure 5A-D, Supplemental Figure 13). These results demonstrate that inhibition of BFT by PIBs is sufficient to ameliorate BFT-mediated pathology *in vivo* and prompted us to pursue oral administration studies.

**Figure 5.**
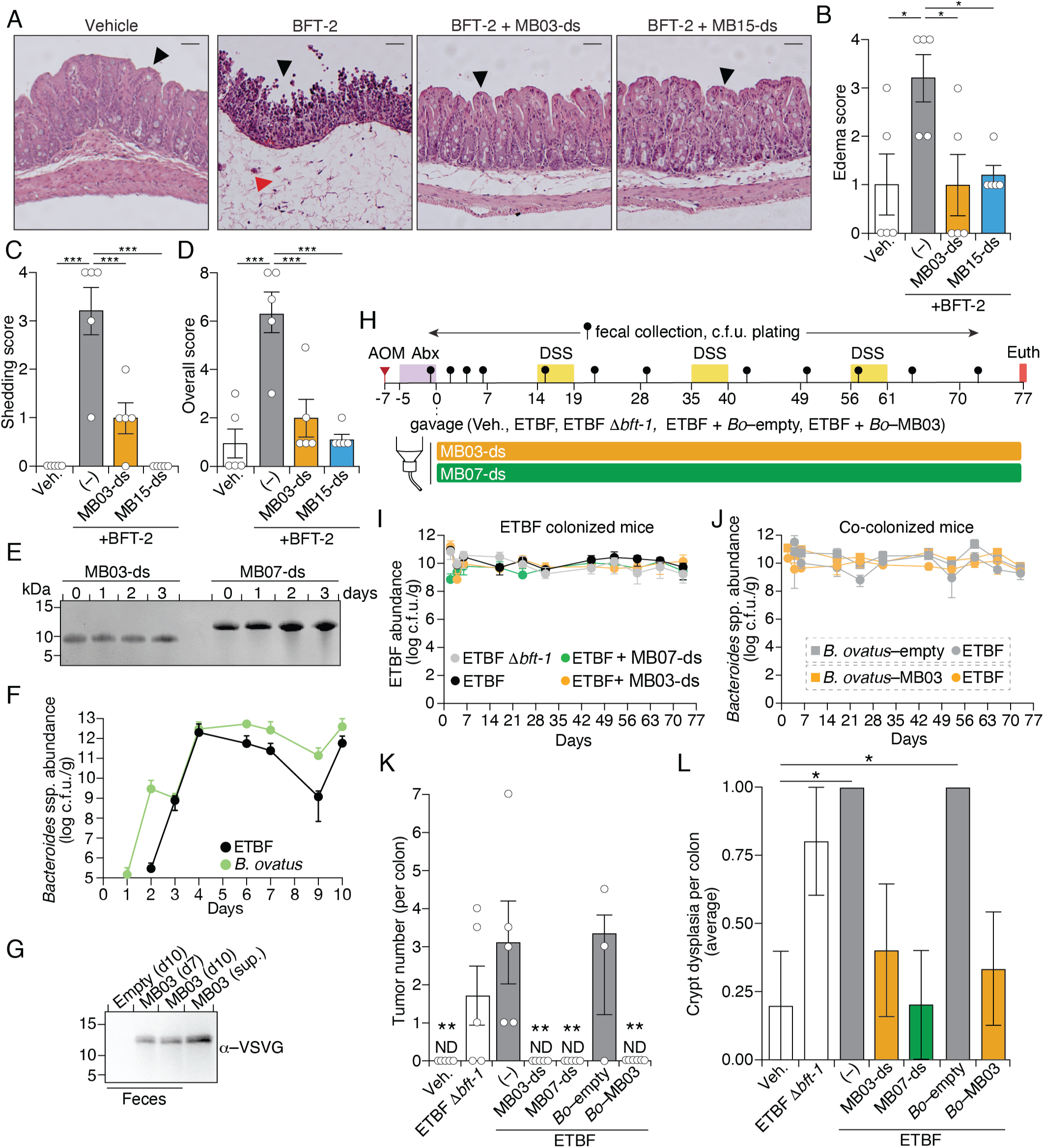
PIBs prevent ETBF pathology in murine models. A) Representative hematoxylin and eosin (H&E)-stained sections of mouse ceca injected with either PBS (vehicle), BFT-2, BFT-2 and MB03-ds, or BFT-2 and MB15-ds. Quantification of cecum histopathology including edema score (B), shedding score (C), and overall score (D). Black arrows indicate intact epithelial crypt in PBS, MB03-ds, and MB15-ds treated mice, with epithelial shedding in BFT-2 treated mice. Red arrow indicates edema formation. E) SDS-PAGE analysis of the indicated minibinders incubated in mouse drinking water. F) Fecal abundance of the indicated strains following their introduction to antibiotic-treated mice via oral gavage on day zero at a 1:1 ration (as determined by OD_600_). Strains were marked with antibiotic resistance cassettes (ETBF, tet; *B. ovatus,* erm) to enable quantification by plating on selective media. G) Western blot analysis of VSV-G–MB03 in fecal pellets collected the indicated number of days post introduction of *B. ovatus–*MB03 via oral gavage. H) Timeline depicting treatments applied and samples collected in an experiment to assess the impact of PIBs on ETBF-induced tumorigenesis in the AOM/DSS-treated mouse model. Mouse colons were harvested and fixed for histological staining at the time of euthanasia. I) Abundance of ETBF (left) or ETBF and *B. ovatus* (right) in AOM/DSS treated mice at the indicated time points post introduction via oral gavage. Antibiotic resistance-marked strains were quantified by plating on selective media as described in (F). K,L) Quantification of tumor counts (K) or crypt dysplasia scores (L) from mice treated with AOM/DSS as depicted in (H) and subjected to the indicated treatments. *Bo, B. ovatus.* MB03-ds and MB07-ds treatment groups indicate mice supplied with MB03-ds or MB07-ds in the drinking water supply. Data in B-D,F,I-L represent means and standard errors. Asterisks indicate values significantly different from the ETBF treatment (B-D, K) or vehicle (L) (1-way ANOVA with Dunnett’s multiple comparisons test (B-D), Kruskal-Wallis test with Dunn’s multiple comparisons test (K), and Fisher’s exact test with Bonferroni correction (L), *p<0.5, **p<0.01, ***p<0.001).

Multiple murine models have been employed to study the inflammatory and tumorigenic effects of BFT^50,69–71^. We opted to employ treatment with the genotoxic carcinogen azoxymethane (AOM) followed by repeated cycles of inflammation-promoting dextran sodium sulfate (DSS), a combination reported to mimic colitis-induced colorectal cancer in humans^72^. Previous studies demonstrate that in this model ETBF colonization exacerbates tumor accumulation in the distal colon^70,73^. We assessed the potential for PIBs to limit ETBF-induced tumorigenesis via two delivery routes: *i*) direct oral administration of disulfide-stabilized PIBs and *ii*) introduction of PIB-secreting LBPs via oral gavage. Direct oral PIB delivery to mice requires the minibinders to be stable in drinking water. To assess this, we incubated MB03-ds and MB07-ds in mouse drinking water for a period of three days (Figure 5E). These PIBs were chosen to interrogate inhibition potency of BFT in *in vitro* assays versus exceptional stability in gastric and intestinal fluids, respectively. SDS-PAGE analysis revealed minimal degradation of either minibinder over this time period, supporting the feasibility of this delivery route (Figure 4H,I).

Successful *in vivo* BFT inhibition by PIB-secreting LBPs requires the strains either stably established in ETBF colonized individuals or regularly supplied. Rather than perform repeated oral gavages, we assessed colonization dynamics of ETBF and *B. ovatus* when co-administered at different initial starting ratios following antibiotic pretreatment (Figure 5F, Supplemental Figure 14A,B). By four days post-gavage, both strains achieved robust colonization levels (10^11^-10^12^ c.f.u./g of feces) in antibiotic pre-treated mice, independent of the initial condition (Supplemental Figure 14B). Colonization levels of both populations persisted through euthanasia at ten days post-gavage. In this experiment we included one mouse colonized with *B. ovatus* secreting MB03 (*B. ovatus*–MB03). By seven days post-gavage, MB03 was readily detected in fecal samples from this mouse, confirming PIB production *in vivo* (Figure 5G).

With evidence supporting the feasibility of our proposed PIB delivery routes, we designed an AOM/DSS-treated mouse study in which we measured ETBF-induced tumorigenesis in mice co-colonized with *B. ovatus*–MB03 or supplied with PIBs in drinking water (Figure 5H). Mice were treated with AOM, and after five days of antibiotic treatment, equivalent levels of ETBF and *B. ovatus*–MB03 (or the control strain *B. ovatus*–empty) were introduced via oral gavage, and weights monitored (Supplemental Figure 15A,B). For those receiving purified PIBs (MB03-ds or MB07-ds), dosage was calculated at a level estimated to supply 2.5 mg/kg daily (Supplemental Figure 15C). Mice were subsequently subjected to three rounds of DSS treatment (1%) and were euthanized on day 77 post-gavage. Analysis of colony forming units from fecal samples collected throughout the experiment indicated stable colonization by ETBF was achieved across groups, with both control and MB03-secreting *B. ovatus* strains achieving similar population densities to ETBF (Figure 5I,J). In mice colonized with *B. ovatus–*MB03, we detected the PIB in fecal samples collected throughout the course of the experiment (Supplemental Figure 15D). Using immunoprecipitation with BFT to enrich for functional PIBs, we also detected MB03-ds in fecal pellets of mice receiving the purified minibinder in drinking water (Supplemental Figure 15E).

The number of tumors present in the distal colon of mice at the conclusion of the experiment was assessed by histological examination by a board-certified veterinary pathologist (Supplemental Table 3). Consistent with previous reports, we found that ETBF colonization correlated with a significant increase in colorectal tumors over control mice, in which no tumors were detected (Figure 5K, Supplemental Figure 15F). Strikingly, we detected no tumors in ETBF colonized mice supplied with MB03-ds or MB07-ds or co-colonized with *B. ovatus* secreting MB03 – but not the *B. ovatus* control strain – consistent with effective inhibition of BFT activity by the minibinders *in vivo* (Figure 5K, Supplemental Figure 15F). Crypt dysplasia scores correlated with tumor counts; average crypt dysplasia per colon was significantly higher in ETBF and ETBF with control *B. ovatus*-colonized mice compared to the PBS control, while mice supplied with PIBs did not exhibit significant dysplasia (Figure 5L). Colonization with ETBF Δ*bft-1* resulted in an intermediate phenotype, with fewer tumors than in the ETBF colonized mice but more than in the control group. We do not have an explanation for this finding, which contradicts previous studies and is inconsistent with the phenotype we observe for mice in which BFT is inhibited by minibinders^74^. Genome sequencing of ETBF Δ*bft-1* and parent strain clones from both the inoculum provided to mice and isolated from fecal samples revealed no genomic differences between the strains outside of the introduced gene deletion, indicating the unexpected phenotype is not due to compensatory mutations in ETBF Δ*bft-1*. Larger *in vivo* studies will be required to more definitively assess the efficacy of PIBs in preventing ETBF-induced colorectal pathology. Nonetheless, our findings provide evidence that orally supplied PIBs hold promise for the prevention of ETBF-linked disease.

## Discussion

In this study, we demonstrate how protein design and engineered live biotherapeutics can be applied to the neutralization of bacterial toxins in the gut. Our work capitalized on two key advantages of designed minibinders over other therapeutic biologics: the potential for their production by LBPs and the readiness by which they can be engineered for enhanced stability. Localized secretion by engineered commensal bacteria, as we demonstrate here, provides a means for sustained delivery of inhibitors directly in the colon, potentially enabling long-term toxin suppression without repeated dosing. In the context of ETBF, where BFT contributes to niche acquisition and microbial competition within the colon, prolonged toxin neutralization may additionally promote pathogen displacement through competitive exclusion by non-toxigenic commensal species^65,75^. Successful application of LBPs in a clinical setting will require strain engraftment, a challenge that has historically limited the realization of the potential of this therapeutic approach^58^. One promising strategy for overcoming this challenge is to supply a privileged nutrient source that the introduced strain – but not the resident microbiota – can metabolize. For instance, provision of seaweed rich in the marine polysaccharide porphyran enabled stable, reproducible establishment of *B. ovatus* and other *Bacteroides* species capable of using this resource^61,76^.

As an alternative to delivery by LBPs that circumvents both engraftment challenges and the potential for regulatory hurdles for their clinical use, we explored the feasibility of direct oral delivery of PIBs. We found that both MB03-ds and MB07-ds successfully prevented ETBF-mediated pathology in a murine model of colorectal cancer, despite the former displaying a relatively short half-life in SGF. We hypothesize that continuous provision of the MBs in the drinking water supply, along with the rapid murine gastric passage time for liquids and small particles (half-emptying time of 15-20 min), enabled sufficiently inhibitory concentrations of the MBs to reach the colon^77,78^. In moving toward development of a clinically effective BFT inhibitor, directed evolution experiments or the introduction of additional disulfide bonds could be applied to further improve PIB stability in SGF and SIF, or strategies could be implemented to improve the binding affinity of PIBs for BFT^33,79^. Encapsulation methods such as the use of enteric coatings to prevent degradation in the acidic gastric environment also hold promise for improving the efficacy of orally administered minibinders^80^. Overall, our work provides support for the feasibility of using orally administered minibinders for therapeutic applications, an application of designed proteins that has to date been minimally explored^33,81^.

Although our strategy for designing BFT inhibitors was initially focused on recapitulating the BFT zymogen aspartate switch to directly occlude the catalytic zinc center, the most effective inhibitors emerged from targeting the prodomain–catalytic domain interface. This finding highlights an advantage of diffusion-based protein design approaches, in that they enable the identification of new binding interfaces. In the case of BFT, inhibition via the prodomain interface fortuitously offers several advantages over an active site-targeting approach. Relative to the catalytic face of BFT, the prodomain-interaction surface is conserved across toxin isoforms, suggesting that PIBs should display broad-spectrum inhibition. This is supported by our findings that PIBs block BFT-2 mediated E-cadherin release from HT-29 cells *in vitro* and prevent tissue damaged associated with direct injection of BFT-2 to the murine cecum. Conversely, when comparing across metzincin-family proteases, the prodomain interface of BFT diverges more sharply from eukaryotic homologs than does the active site, suggesting PIBSs pose a lower risk than active site binders for off-target inhibition of related eukaryotic enzymes^50^. Additionally, we observed that PIBs inhibit BFT activity through both blocking the interaction with the receptor claudin-4 and by inhibiting its proteolytic activity in the absence of receptor binding. The mechanism by which PIBs directly inhibit BFT activity is unclear. AF models suggest that PIB binding has minimal impact on the BFT active site confirmation, but subtler allosteric effects such as limiting the mobility of active site residues or destabilizing the zinc binding site could explain their inhibitory effects. More broadly, our work supports the notion that regulatory and receptor binding interfaces of toxins represent particularly attractive surfaces for inhibitor development. Indeed, inhibitors designed to act as decoys that mimic toxin receptors have been developed for several toxins, and minibinders designed to target the receptor binding interface of *C. difficile* toxin B exhibit broad spectrum activity towards toxin isoforms^44,79,82^.

While our *in vivo* studies largely focused on testing the efficacy of PIBs in preventing ETBF-associated tumorigenesis, epidemiological studies suggest that this organism is an important agent of inflammatory gastrointestinal diseases more broadly. Further development of PIBs as therapeutic agents could enable treatment of a range of ETBF-associated pathologies, such as inflammatory diarrhea and Crohn’s disease^15,16,83,84^. Additionally, PIBs could find utility as a research tool to gain greater understanding of the mechanisms by which ETBF causes disease, which remains poorly understood. Our finding that PIBs limit BFT activity by blocking claudin-4 binding supports the physiological relevance of the BFT-claudin-4 interaction. Moving forward, future studies should further address the therapeutic potential of PIBs and explore routes to regulatory approval for their use in treating ETBF-mediated disease.

## Methods

### Minibinder design

#### Minibinders targeting the BFT-1 catalytic site

Minibinder scaffolds were designed against the catalytic domain of BFT-1 using a published crystal structure (PDB ID: 8H3Y, chain A, residues 212-397)^37^. Catalytic hotspot residues 348/349/352/358/368 were specified based on prior functional studies^37^. De novo backbone generation was performed using a denoising diffusion framework^32^, yielding 10,000 miniprotein backbones positioned proximal to the catalytic site. Sequence design was performed using ProteinMPNN^39^, generating 20,000 candidate minibinder sequences. Structural validation of an interaction between designs and BFT was carried out using AF2^40^, and designs were evaluated using interchain Predicted Aligned Error (iPAE) and predicted Local Distance Difference Test (pLDDT). Additional filtering metrics, including spatial aggregation propensity (SAP) and binding free energy ΔΔG, were calculated using Rosetta^38^. Designs meeting the following thresholds were retained: iPAE < 6, pLDDT > 80, SAP < 35, and Rosetta ΔΔG < -30. This yielded 146 high-confidence designs. From these 146 designs, one design was predicted to bind at an alternative surface corresponding to the prodomain interface, indicating the presence of an energetically favorable binding site outside the predefined catalytic hotspot region.

To further diversify the design space, 140 candidates were subjected to partial diffusion, generating 7,000 additional backbone conformations (50 partial diffusion trajectories per input structure). Two sequence design strategies were subsequently applied. In the first approach, LigandMPNN, which explicitly models coordination with the catalytic zinc ion, was used to generate one sequence per backbone (7000 total)^42^. AF2 evaluation identified 6,003 designs with favorable predicted interactions. In parallel, ProteinMPNN was applied without explicit modeling of the zinc ion, generating four sequences per backbone (28,000 total)^39^. Of these, 8,727 designs exhibited high-confidence binding (PAE interaction score < 4.9). 14,730 designs top-ranked across both approaches were selected for downstream screening via yeast surface display.

#### Minibinder design to the BFT-1 prodomain interface

##### 45-65 amino acid designs

De novo diffusion was used to generate 10,000 backbone scaffolds targeting hotspot residues 234/259-261/265^32^. Sequence design with ProteinMPNN yielded 100,000 candidate minibinders^39^. Designs were evaluated using AF2 and Rosetta-based metrics. Filtering criteria (PAE interaction ≤ 10, contact molecular surface > 300 Å^2^, ΔΔG < -20, SAP score < 35, and binder Cα r.m.s.d. < 1.5 Å) resulted in 1,739 high-confidence candidates. To further explore the local structural space, partial diffusion was applied (50 trajectories per input), generating 86,950 additional backbone designs. Sequence optimization using ProteinMPNN (four sequences per backbone) yielded 347,800 candidates. AF2 predictions were used to evaluate binding confidence and structural quality. Designs were filtered using stringent criteria (PAE interaction ≤ 6, contact molecular surface > 300 Å^2^, Rosetta ΔΔG < -48, SAP score < 35, and binder Cα r.m.s.d. < 1.5 Å), resulting in 12,402 candidates for yeast display screening.

##### 66-88 amino acid designs

A complementary design campaign targeting longer minibinders (66-88 amino acids) was performed using the same prodomain interface. De novo diffusion generated 9,877 backbone scaffolds, and ProteinMPNN produced 10 sequences per scaffold. Following AF2 and Rosetta-based evaluation, 844 high-confidence designs were retained. Partial diffusion further expanded this set, generating 42,200 additional backbone designs. Sequence optimization (four sequences per backbone) was followed by AF2 evaluation and filtering (PAE interaction ≤ 9, contact molecular surface > 300, ΔΔG < -33, SAP score < 35, and binder Cα r.m.s.d. < 1.5 Å), yielding 11,843 candidates for experimental screening.

### Strains and culture conditions

All strains used in this study are listed in Supplemental Table 4. *Escherichia coli* strains EC100D λ pir+ and DH5α were used for cloning, S17-1 λpir for conjugation, and BL21(DE3) or Rosetta(DE3) for protein expression. *Bacteroides* strains used include enterotoxigenic *B. fragilis* VPI-13784 (BFT-1) and 86-5443-2-2 (BFT-2), nontoxigenic *B. fragilis* NCTC 9343, and *B. ovatus* ATCC 8483. *Saccharomyces cerevisiae* strain EBY100 was used for yeast display. *E. coli* strains were grown aerobically at 37 °C in lysogeny broth (LB). *Bacteroides* strains were cultured at 37 °C under anaerobic conditions in pre-reduced brain heart infusion (BHI) supplemented with 0.5 g L^-1^ cysteine, 0.01 ng mL^-1^ Vitamin K1, and 50 μg mL^−1^ hemin (BHI-S). Anaerobic culturing was performed in an anerobic chamber (Don Whitley Scientific) containing 80% N_2_, 10% CO_2_, and 10% hydrogen. Yeast were grown in selective minimal medium (details, ref(-ura, -trp))^85^. Antibiotics and supplements were added to media as needed at the following concentrations: 200 μg mL^−1^ gentamicin (gent), 5 or 25 μg mL^−1^ erythromycin (erm), 5 μg mL^−1^ tetracycline (tet), 150 μg mL^−1^ carbenicillin (carb), 25 μg mL^−1^ kanamycin (kan), 34 μg mL^−1^ chloramphenicol (cam), 100 ng mL^−1^ anhydrotetracycline (aTC), and 5% defibrinated horse blood (Lampire Biological Products or Quad Five).

### Yeast display and fluorescence activated cell sorting

DNA libraries encoding minibinder designs were synthesized commercially (Agilent). Libraries were cloned into pETCON for surface display as Aga2p fusions with a C-terminal Myc tag^31,86,87^. Yeast strain EBY100 was transformed by electroporation and grown in selective minimal media, and expression was induced with 2% galactose. To prepare libraries for sorting, yeast cells were spun down 16-24 hours post-induction, washed with PBS supplemented with 1% bovine serum albumin, and incubated for 30-60 minutes with BFT-1–H_8_ and antibodies to detect minibinder surface expression (α-Myc-FITC conjugate, Immunology Consultants Laboratory) and BFT-1–H_8_ (α–His-PE conjugate, Bio-Techne), both added at a 1:100 dilution. The stained cells were then sorted by fluorescence-activated cell sorting (FACS) with sequential gating selected for intact cells, singlets, minibinder expressing populations, and high-affinity binders (top 1-5% of the population). Libraries were enriched over two rounds of sorting followed by sorts with progressively decreasing BFT-1–H_8_ concentrations (Library 1: 1 μM to 1 nM; Library 2: 1 μM to 100 nM).

### Plasmid construction

All plasmids, primers, and constructs are listed in Supplemental Table 4. DNA fragments were obtained from Integrated DNA Technologies (IDT) or from Twist Bioscience. Constructs were assembled using Gibson assembly or Golden Gate cloning and verified by Sanger sequencing. Allelic exchange constructs for *Bacteroides* using the pLGB13 vector were generated by amplifying ∼1 kb homology arms from isolated genomic DNA (DNeasy Blood & Tissue kit, Qiagen)^64^. Constructs for protein expression or antibiotic resistance cassette insertion in *Bacteroides* species were assembled in pNBU2-based vectors with codon optimized sequences^88^.

### Generation of genetically modified Bacteroides strains

Constructs for allelic replacement or ectopic gene insertion were delivered to *Bacteroides* strains via triparental matings as previously described^64^. Briefly, *E. coli* S17-1 λ pir strains containing either pLGB13 or pNBU2 derivatives were mixed together with *E. coli* S17 carrying the helper plasmid RK213 with a ten-fold excess of the target recipient *Bacteroides* strain and incubated aerobically overnight at 37°C. For allelic exchange using pLGB13, merodiploids were selected by plating on media containing gent and erm, then grown in non-selective liquid medium to promote plasmid excision and counter-selected by plating on aTC supplemented media. Colonies were patched onto erm containing plates to confirm plasmid loss. Mutations were confirmed by PCR and Sanger sequencing (Genewiz). For pNBU2 matings, plasmids were mobilized as described above and selected on antibiotic-containing media. Insertion was confirmed by PCR and Sanger sequencing.

### BFT-1–H_8_ purification

One liter of degassed BHI-S was inoculated with *B. fragilis* VPI 13784 *bft–*H_8_ scraped from a freshly grown, confluent agar plate. Cultures were grown anaerobically for 16 hours (OD_600_ of between 1.8-2.1). At harvest, 5× buffer (2.5 M NaCl, 250 mM Tris-HCl, pH 7.5, 25 mM imidazole) was added and cultures centrifuged at 10k × *g* for 30 minutes. Supernatant was collected, filtered (0.22 µm PES membrane) and loaded onto an AKTA fast protein liquid chromatographer (FPLC) using a HisTrap HP column (Cytiva). Bound protein was eluted using a linear imidazole gradient from 10 to 250 mM. Protein purity was assessed by sodium dodecyl sulfate-polyacrylamide gel electrophoresis (SDS-PAGE) and silver staining. Fractions with high purity were concentrated using a 10 kDa molecular weight concentrator (Amicon), then further purified using a HiLoad 16/600 Superdex 200 pg column (GE Healthcare) equilibrated with sizing buffer (200 mM NaCl, 50 mM Tris-HCl pH 7.5 and 5% glycerol (v/v)). Concentrated fractions with high purity were aliquoted, flash frozen in liquid nitrogen, and stored at -80 °C until use.

### BFT-2–H_6_ purification

Recombinant proBFT (residues D24-D397) with a C-terminal 6×His tag was expressed from a modified pET30a vector in *Escherichia coli* Rosetta(DE3). Overnight cultures of *E. coli* were subcultured at a 1:50 ratio into LB with kan and cam, grown at 37 °C and 240 rpm until they reached an optical density (OD)_600_ of 0.4–0.6 and then induced with 0.5 mM isopropyl β-D-1-thiogalactopyranoside (IPTG) at room temperature overnight. Cells were collected by centrifugation at 3,000 ×*g* for 15 min at 4 °C and resuspended in buffer A (50 mM Tris-HCl, 500 mM NaCl, 20 mM imidazole, pH 7.5). Cells were lysed by sonication and cellular debris removed by centrifugation at 17,200 × *g* for 45 min at 4 °C. Lysates were filtered through a 0.22 µm filter and loaded onto a HisTrap column (Cytiva). Protein was eluted with five CV of buffer A containing 250 mM imidazole. Fractions containing the A_280_ peak were pooled and dialyzed into PBS overnight. The resulting proBFT was treated with trypsin (Sigma) at a 1:30 w/w ratio (trypsin:proBFT) to cleave off the BFT N-terminal prodomain, repurified on a HisTrap column, and dialyzed into PBS overnight. Purity was assessed by SDS-PAGE followed by Coomassie staining, and fractions with high purity were concentrated, aliquoted, flash frozen in liquid nitrogen, and stored at -80 °C until use.

### Untagged BFT purification

Recombinant proBFT (residues D24-D397) with an N-terminal 6×His tag was expressed from a modified pET28a vector in *Escherichia coli* Rosetta(DE3). Cells were grown and induced as described above. Cells were collected by centrifugation at 4,000 ×*g* for 20 min at 4 °C and resuspended in buffer A. Cells were lysed using a Cell Disruptor (Constant Systems) and cellular debris removed by centrifugation at 20k × *g* for 40 min at 4 °C. Clarified lysate was incubated with HisPur Ni-NTA resin (ThermoFisher) for one hour at 4 °C with stirring. The mixture was loaded onto a gravity column. Resin was washed with ten CV of buffer A and eluted with 5 CV of buffer A with 300 mM imidazole. Eluate was buffer exchanged into 50 mM Tris-HCl, 150 mM NaCl, pH 7.5 using a PD-10 desalting column (Cytiva) then trypsin treated as described above, resulting in release of the catalytic domain of BFT from the 6×His tagged N-terminal prodomain. The reaction was stopped using 1 mM PMSF and buffer exchanged into 20 mM Tris-HCl, 20 mM NaCl, pH 7.5 using an ultrafiltration device. Sample was injected onto a Mono Q 5/50 GL column (Cytiva) and mature BFT was eluted using a gradient from 20 to 500 mM NaCl. Purity was assessed by SDS-PAGE followed by Coomassie blue staining. Concentrated fractions with high purity were aliquoted, flash frozen in liquid nitrogen, and stored at -80 °C until use.

### Minibinder purification

Overnight cultures of *E. coli* BL21(DE3) carrying pET28+ constructs expressing minibinders and pCDFDuet-BirA for *in vivo* biotinylation when appropriate were back diluted 1:100 in LB and grown at 37 °C with shaking at 200 r.p.m. until the OD_600_ reached 0.4-0.8. The incubation temperature was then reduced to 18 °C, and samples were induced with IPTG (0.1 mM final) and grown overnight. For Avi-tagged binders, biotin (60 µM final) was also added at the induction step to promote biotinylation. Cells were harvested by centrifugation at 6000 × *g* for 30 minutes and resuspended in lysis buffer (500 mM NaCl, 50 mM Tris-HCl, pH 7.5, 5 mM imidazole, 0.1% Triton-X100, 0.5 mg mL^-1^ lysozyme, and 1 mU benzonase). Cells were lysed by sonication and cellular debris removed by centrifugation at 20,000 × *g* for 60 min at 4 °C. Lysates were filtered through a 0.22 µm filter then purified using their respective affinity tag (H_6_ or biotinylated Avi). H_6_-tagged minibinders were purified using a 1 mL HisTrap FF column as described above. Fractions with high purity confirmed via SDS-PAGE were concentrated using a 3 kDa molecular weight cutoff (Amicon) and further purified on a Superose 6 Increase 10/300 GL column equilibrated with sizing buffer (500 mM NaCl, 50 mM Tris-HCl pH 7.5). For biotinylated minibinders, clarified lysate was loaded to a gravity column packed with streptavidin mutein matrix (Roche), and purified per manufacturer’s instructions. Protein purity was assessed as above, and selected fractions further purified by size-exclusion chromatography as for H_6_–tagged minibinders.

### Surface plasmon resonance assays

Binding affinities were measured using single cycle kinetic experiments on a Biacore 8K (Cytiva)^89^. Measurements were obtained by immobilization of 1 µM of purified BFT-1–H_8_ protein on an NTA chip (Cytiva), and minibinders were injected in a nine-point concentration series, with an association time of 90 seconds and dissociation time of 300 seconds. Sensorgrams were double-referenced and fitted with a 1:1 binding kinetic fit model using Biacore Insight evaluation software.

### Protein precipitation assays

Direct interactions between BFT-1–H_8_ and minibinders were probed using avi-tagged minibinders following previously published protocols with minor modifications^90^. One µM of BFT-1– H_8_ was combined with 200 nM biotinylated minibinder in buffer (500 mM NaCl, 50 mM Tris-HCl, pH 7.5, 10 mM imidazole, 0.01% Tween-20). Protein mixtures were incubated with 50 μL of Ni-NTA resin (Qiagen) at RT for 30 min with rotation. Agarose beads were pelleted and washed five times with wash buffer (500 mM NaCl, 50 mM Tris-HCl, pH 7.5, 10 mM imidazole). Bound proteins were eluted with elution buffer (500 mM NaCl, 50 mM Tris-HCl and 500 mM imidazole), mixed with 4× Laemmli loading buffer (Bio-Rad), boiled 10 min at 95 °C, and subjected to Western blot analysis.

### Western blot analysis

Proteins were resolved by SDS-PAGE (8-16% acrylamide gels, Bio-Rad). For minibinder and BFT-1–H_8_ detection, proteins were transferred to nitrocellulose membranes at 100 V for 30 min at 4 °C. For detection of RpoB (control intracellular protein used in secretion assays), proteins were transferred to a PVDF membrane for 7 minutes (20 V for 1 min, 23 V for 4 min, 25 V for 2 min) using an iBlot2 (Invitrogen). Membranes were blocked in 5% bovine serine albumin (BSA) in Tris-buffered saline with Tween 20 (TBS-T) for 1 hour at room temperature. Blots were incubated in primary antibodies at room temperature for one hour (1:5000 in 5% BSA for α–His HRP-conjugated (Qiagen) and α–VSV-G (Millipore Sigma or Novus Biologicals), 1:5000 in 1% BSA for streptavidin-HRP conjugated (Millipore), and 1:10000 dilution in 3% BSA for α−RNA polymerase B (Biolegend)). Blots were washed four times with TBS-T. α−VSV-G and α−RpoB blots were incubated with secondary antibody for one hour at room temperature (1:5000 α−rabbit HRP-conjugated (Millipore Sigma), 1:5000 α−mouse HRP-conjugated (EMD Millipore), respectively), then washed four times with TBS-T. Blots were developed using Clarity Max Western ECL Substrate (Bio-Rad) and visualized using an Invitrogen iBright 1500 imager.

For the BFT-2– H_6_ Western blots, proteins were transferred to nitrocellulose membranes at 100 V for 1 hour in methanol-supplemented NuPAGE transfer buffer (Invitrogen). Blots were blocked in 5-10% dry milk in PBS-T (0.05% Tween-20 in PBS, pH 7.4) for one hour. Primary antibodies were diluted in blocking buffer and incubated with membranes for one hour. Primary antibodies include α−β-actin (1:2,000, Sigma), α−His (5 μg ml^-1^, R&D Systems), and α−E-cadherin ectodomain (1:250, Invitrogen). Membranes were washed three times with PBS-T then incubated for one hour with HRP-conjugated secondary antibodies diluted in blocking buffer (α−mouse, 1:1000, Kindle Biosciences; α−rat, 1:1000, Cell Signaling Technology). Following three washes in PBS-T, blots were developed using SuperSignal West Pico PLUS chemiluminescent substrate (Thermo Fisher) and imaged on an iBright CL1500 (Invitrogen).

### Bio-layer interferometry

Binding interactions were measured on an Octet R4 (Sartorius) at 25 °C with orbital shaking at 1000 rpm. Streptavidin (SA)-coated biosensors were hydrated for at least 10 minutes in assay buffer (25 mM tris, pH 7.5, 150 mM NaCl, supplemented with 0.01% Tween-20 and 0.1% BSA). MB03 variants were loaded onto biosensors at 100 nM to achieve a loading response of ∼1-1.5 nm. Following loading, biosensors were equilibrated in fresh assay buffer for 120 seconds to establish a baseline. Association kinetics were measured by transferring biosensors into wells containing a titration series of BFT-1–H_8_ until equilibrium was approached. Dissociation was then monitored by returning biosensors to fresh assay buffer. Steady-state and global kinetic analyses were performed using the manufacturer’s software (Octet Analysis Studio 13.0) with a 1:1 binding model.

### Human cell culture

HT-29 (ATCC HTB38), HT-29 claudin-4 KO, and HT-29-MTX-E12s cells (Sigma, 12040401- 1VL) were maintained at 37 °C in 5% CO_2_ in DMEM (Corning, #10-013-CV) supplemented with 10% FBS (Fisherbrand, #FB12999102), 1× NEAA (Corning, #25-025-CI), and 1× pen/strep (Corning, #30-002-CI). Cell stocks were passaged weekly and cultured in T-25 flasks (2 × 10^4^ cells mL^-1^). HT-29 cells were used between passage numbers 136-148 and HT-29-MTX-E12s were used between passage numbers 51-64. All cells were tested and confirmed to be negative for mycoplasma.

### BFT-inhibition assays

HT-29 cells were seeded onto a 96-well plate at 2.5×10^4^ cells mL^-1^ and grown to reach 60-75% confluency. Supernatant from *B. fragilis* culture overnights (*B. fragilis* VPI-13784 for BFT-1) was collected by centrifugation, followed by filtration (0.22 µm filter). Supernatants were applied at a 0.25 dilution to each well. Normalized volumes of inhibitor stocks were applied to wells, and samples incubated aerobically at 37 °C overnight. Cell supernatants were then analyzed via ELISA according to manufacturer’s protocols (E-cadherin #DY648 and IL-8 #DY20805, R&D Systems). Cell images were taken of live, unstained cells pre- and post-incubation using the Cytation 5 (Agilent) as a quality control measure for cell integrity. Adherent cells post-imaging were used for MTT viability assays using a CyQUANT MTT kit (Invitrogen) according to the manufacturer’s rapid protocol.

### Cellular BFT binding assays

WT claudin-4 or claudin-4 KO HT-29 cells seeded in 12-well plates were rinsed once with serum-free culture medium, treated with H_6_–BFT-2 (100 ng mL^-1^, ∼5 nM) in serum-free cell culture medium at 37 °C and 5% CO_2_ for 40 minutes and rinsed twice with ice-cold PBS to remove unbound BFT-2. Cells were then lysed directly in the plate for detection in SDS loading buffer (50 mM Tris-HCl, 2% SDS, 10% glycerol, 0.02% bromophenol blue, 100 mM DTT, pH 6.8). Cells were solubilized with orbital shaking for five min at 4 °C, heating at 100 °C for ten minutes, and then debris was pelleted by centrifugation at 17,200 × *g* for five minutes. Clarified lysates were stored as single-use aliquots until used for immunoblotting. Samples were loaded onto NuPAGE 4-12% Bis-Tris Miniprotein gels (Invitrogen), and electrophoresis was performed at 200 V for 35 min in NuPAGE MES SDS running buffer (Invitrogen) and analyzed via Western blot^50^.

### Protein cleavage assays

Purified BFT cleavage activity on monomeric G-actin was assessed under varying pH (6-9), temperature (RT to 55 °C), and calcium concentration (1 mM CaCl_2_ or no calcium added)^91,92^. G-actin was prepared according to manufacturer’s instructions (Cytoskeleton Inc). Actin (2 µM final concentration) was incubated with purified BFT at a 100:1 substrate-to-enzyme molar ratio under the indicated conditions in cleavage buffer (25 mM Tris-HCl, 150 mM NaCl, and 2 µM Zn_2_Cl). Reactions were quenched at 30 minutes and 5 hours with 2× Laemmli buffer (Bio-Rad) followed by boiling at 95 °C for 10 minutes. Samples were resolved by 6-18% SDS-PAGE (Bio-Rad) at 120 V for 60 minutes and stained with Coomassie blue.

Optimal enzymatic conditions (pH 7, 37 °C incubation) were used for subsequent assays. Under these conditions, G-actin or E-cadherin (R&D Systems) were incubated with BFT-1 at concentrations described above in the presence or absence of 1 µM MB03. Trypsin served as a positive control (Sigma-Aldrich). Reactions were processed and analyzed as described above.

### Bacteroides protein secretion assays

*B. ovatus* and non-toxigenic *B. fragilis* strains engineered to secreted minibinders (or vector only control strains) were inoculated into pre-reduced BHI-S and anaerobically grown for 16 hours at 37 °C. Cultures were centrifuged at 8,000 × *g* for 30 min to pellet cells and collected supernatant filtered (0.22 µm filter). Cell pellets normalized by total protein concentration were resuspended in 2× Laemmli loading buffer (Bio-Rad). Normalized supernatant volumes were concentrated 50-fold via trichloroacetic acid precipitation^93^ and resuspended in 2× Laemmli loading buffer (Bio-Rad). Samples were boiled for 10 minutes, and subject to Western blot analysis.

### Live bacteria BFT-inhibitory assays

Turbid overnight cultures of *B. fragilis* or *B. ovatus* cultures were resuspended in DMEM and normalized to an OD = 0.3 for *B. fragilis* or OD = 6.4 for *B. ovatus* for a 1:10 ratio of *B. fragilis* to *B. ovatus*. *B. ovatus* strains were serially diluted to achieve a 1:1 or 10:1 ratio while maintaining a constant level of *B. fragilis*. All bacteria were applied at an equal volume to HT-29 cells prepared as above. Samples were incubated anaerobically for five hours at 37 °C, and cell supernatants collected for ELISAs and processed as described above.

### Tissue culture microscopy

HT-29 (p143) cells were seeded onto an 8-well chamber slide (Falcon #354118) 11.6×10^4^ cells mL^-1^ and to reach ∼60-75% confluency. Minibinders were applied at 1 µM, purified BFT-1–H_8_ was applied at 100 nM, and ETBF SCS was applied as a 0.25 dilution and incubated aerobically for 16 hours at 37 °C with 5% CO_2_. Wells were fixed with 100% cold methanol (Fisher, A413-500) and stained with 1:20 Giemsa solution (Fisher, SDGS500). Well boundaries were removed from the chamber slides and washed with PBS. Cell rounding images were collected using a Leica DM750 with an attached Leica ICC50 HD camera. Images were collected using the 40× objective lenses and processed using the LAS EZ software (vers 3.4.0). Scale bar was determined by imaging a Hausser Scientific (#3110V) hemacytometer at the same magnification as the cell images.

### Colonic organoid assays

Procedures for harvesting mouse colonic organoids, or colonoids (p2) derived from C57BL/6 mouse carcasses was adapted from previously published protocols^94^. Briefly, the colon was dissected, cut longitudinally, washed with PBS + 0.04% bleach, then incubated in dissociation media (PBS, 0.5 mM DTT, 2 mM EDTA). Colon tissue was transferred to PBS and crypt fractions collected. Fractions were counted, pooled, centrifuged, and resuspended in Matrigel (Corning). Mouse colonic crypts were seeded at 3000-5000 crypts per well (20 μL Matrigel in 24-well plates) and maintained for three days in complete media containing DMEM/F12 (Gibco) supplemented with 10% FBS, 0.5× GlutaMax (Fisher), 5 mM HEPES, 0.5× N2 Supplement, 0.5× B27 Supplement, 50 ng mL^-1^ mEGF, and 50% conditioned media from L-WRN cells (ATCC CRL-3276) which produce Wnt3a, R-spondin 3, and Noggin. Cells were passaged via mechanical shearing and maintained for three to five days before freezing in advanced DMEM/F12 (Gibco) supplemented with 25% FBS, 0.5× pen/strep, 0.5× GlutaMax (Fisher), 5 mM HEPES, and 10% DMSO. One week prior to assays, organoids were thawed and maintained in 4 μL of Matrigel in 96-well plates with serum-free media at 37 °C with 5% CO_2_.

For assays, wells were treated with 100 nM BFT-1–H_8_, 10 µM purified minibinder, or vehicle control. Images were collected using the Cytation 5 set to capture a 12-region montage with 8-slice z-stack per well region. Composite images for each well were created using a custom macro in ImageJ to flatten the z-stacks and stitch together the montage. Blinded phenotypic counts were conducted and averaged across two individuals.

### Transepithelial resistance assays

HT-29-E12-MTX cells (p59-60) were seeded onto 24-well transwell inserts (3 µm pore) at 1×10^5^ cells mL^-1^ and grown to form monolayers or until the transepithelial electrical resistance (TEER) ≥200 Ω cm^-2^ (two to three weeks). Wells and transwell inserts were washed twice with PBS, then refreshed with DMEM. Bacterial cultures were normalized to OD = 1.5 (*B. fragilis* VPI-13784) or OD = 2.5 (*B. ovatus* ATCC 8483) in DMEM. Bacterial treatments were pre-mixed as needed prior to application to the apical chamber and incubated anaerobically for five hours at 37 °C with shaking (Belly Button orbital shaker, IBI Scientific, BBULS0001). TEER was measured before and after the incubation using an ERS-2 voltohmmeter (Sigma).

### Minibinder stability assays

Simulated gastric fluid (SGF) was prepared by adjusting 45 mM of NaCl in water with 100 mL HCl to a final pH of 2^95^. Pepsin was added to 2 mg mL^-1^ immediately before use. Simulated intestinal fluid (SIF) was prepared by dissolving 19 mM maleic acid, 34.8 mM NaOH, and 68.6 mM NaCl. Sodium taurocholate (3 mM) and lecithin (0.2 mM) were added, and the pH was adjusted to 6.5^79^. Trypsin and chymotrypsin at 0.1 mg mL^-1^ were added immediately before use.

Minibinders were added to either SIF or SGF at 25 µM and incubated at 37 °C. Reactions were analyzed at 0 min, 15 min, 1 h, 4 h, and 24 h. SGF incubations were neutralized to pH 7 with 0.2 M NaOH. SIF reactions were quenched with PMSF. Samples were boiled for 10 min before analysis via SDS-PAGE. To test stability in water, minibinders were added directly to water provided by the animal facility and incubated at room temperature before analysis via SDS-PAGE.

### X-ray crystallography

All crystallization experiments were conducted using the sitting drop vapor diffusion method. Crystallization trials were set up in 200 nL drops using the 96-well plate format at 20 °C using a Mosquito LCP (SPT Labtech), then imaged using a UVEX PS-256 imager and UVEX microscopes (JAN Scientific). Diffraction quality crystals formed in 24 % PEG 1500 (w/v) and 20% glycerol (w/v) for MB03-ds; 0.2 M lithium sulfate, 0.1 M sodium acetate pH 4.5, and 50 % PEG 400 (w/v) for MB07; 0.1 M citric acid pH 3.5, 2.0 M ammonium sulfate for MB07-ds.

Diffraction data was collected at the Advanced Photon Source on beamline 24-ID-E and the National Synchrotron Light Source II on beamline 17-ID-1 (AMX). X-ray intensities and data reduction were evaluated and integrated using XDS and merged and scaled using POINTLESS and Aimless in the CCP4 program suite^96,97^. Structural determination and refinement starting phases were obtained by molecular replacement using Phaser and the AlphaFold3 (AF3) models for each structure^98^. Following molecular replacement, structures were improved using phenix.autobuild with simulated annealing, and further refined in Phenix^99^.

Model building was performed using COOT^100^. Model quality was evaluated using MolProbity^101^. X-ray structure data collection and refinement statistics are recorded in Supplemental Table 2. Data deposition, atomic coordinates, and structure factors reported in this paper have been deposited in the Protein Data Bank (PDB) with accession codes 36LX, 36LY, and 36MB.

### BFT cecal injection assay

All animal experiments were approved by the Institutional Animal Care and Use Committees of Boston Children’s Hospital. For BFT cecal injection experiments, untagged BFT-2 (10 μg), MB03-ds (60 μg) or MB15-ds (60 μg) was incubated either alone or together in 200 μL of PBS for 30 min at room temperature. Five-week-old CD-1 outbred female mice (Envigo, 030-US) were fasted for 12 hours before injection, anaesthetized with isoflurane and the cecum was exposed via laparotomy. The BFT, PIBs, or BFT:PIB mixture was injected directly into the cecum, followed by wound closure with stitches. Buprenorphine (0.1 mg kg^−1^) was injected subcutaneously immediately after surgery for analgesia. Mice were allowed to recover, and 4 h after surgery, the mice were euthanized. The cecum was collected, rinsed with PBS to remove the contents and fixed with 10% formalin overnight for further histology and immunofluorescence analysis.

The fixed cecum was dehydrated with 70% ethanol, 96% ethanol, 100% ethanol, then xylene (each step twice for 30 min), then infiltrated with melted paraffin at 58 °C for 45 min and embedded into the block. The tissue blocks were sectioned into 6-μm-thick sections. For hematoxylin and eosin (H&E) staining, sections were heated at 42 °C for 30 min and treated with xylene (twice for 5 min), 100% ethanol (thrice for 1 min), tap water (1 min), Gill 3 hematoxylin (2 min), tap water (2 min), eosin (3 min), 100% ethanol (thrice for 1 min) and xylene again (twice for 2 min). Sections were mounted with DPX mountant (Sigma) for histology analysis. Stained sections were coded and scored blindly.

### AOM/DSS-induced colorectal tumorigenesis assays

For pilot colonization studies, four-week-old male C57BL/6J mice (Jackson Laboratory) received streptomycin (5 g L^-1^) and clindamycin (100 mg L^-1^) in drinking water for five days. Studies were performed with three mice per group. Clearance of the native gut microbiome was confirmed by anerobic culture of fecal samples on non-selective media prior to bacterial administration. Bacterial strains were cultured anaerobically in BHI-S medium and prepared for oral gavage in sterile PBS. ETBF^Tet^ VPI-13784 and ETBF^Tet^ Δ*bft-1* VPI-13784 were administered at 10^8^ colony forming units (c.f.u) per mouse. For colonization optimization experiments, *B. ovatus*^Erm^-empty ATCC 8483 or *B. ovatus*^Erm^-MB03 ATCC 8483 were co-administered with ETBF at ratios ranging from 1:10 to 100:1 based on OD_600_-derived c.f.u. estimates. Bacteria were introduced to mice via oral gavage (100 μL per mouse).

Fecal pellets were collected at regular intervals to monitor colonization. Samples were weighed, homogenized in PBS, serially diluted, and plated on selective media containing the appropriate antibiotics. Plates were incubated anaerobically at 37 °C and colony counts were normalized to fecal mass. Remaining material was stored at -80 °C for downstream analyses.

For disease studies, six- to eight-week-old male C57BL/6J mice received a single intraperitoneal injection of 10 mg kg^-1^ azoxymethane (Sigma-Aldrich). Two days later, mice were treated with streptomycin (5 g L^-1^) and clindamycin (100 mg L^-1^) in drinking water for five days, followed by oral gavage with bacterial strains. ETBF^Tet^ and ETBF^Tet^ Δ*bft-1* was administered at 10^8^ c.f.u. *B. ovatus*^Erm^-empty or *B. ovatus*^Erm^-MB03 were co-administered at 10^8^ c.f.u. Mice receiving oral minibinder treatment at 0.015 mg mL^-1^ after gavage obtained an average dose of approximately 2.5 mg kg^-1^ to 3.0 mg kg^-1^ of either MB03-ds or MB07-ds for the remaining duration of the experiment.

Two weeks after bacterial administration, mice underwent three cycles of dextran sodium sulfate (DSS, 1%) treatment (36–50 kDa; MP Biomedicals, Santa Ana, CA, USA) for five days followed by a 16-day recovery period with untreated water. Body weight, fecal samples, and bacterial colonization were monitored weekly. Following the last cycle, mice were euthanized, blood and colon tissues were collected. Serum was isolated from coagulated blood according to published protocols^102^. Colons were flushed with PBS, opened longitudinally, Swiss-rolled, and fixed in 10% formalin. Fixed tissues were processed for H&E staining by the Yale University Comparative Pathology Research Core. Sample sizes for each experiment are reported in the corresponding figure legends. Histological evaluation and scoring were performed by a board-certified veterinary pathologist blinded to treatment groups.

ETBF^Tet^ VPI-13784 and ETBF^Tet^ Δ*bft-1* VPI-13784 isolates were recovered from the fecal samples collected on day 70 and were subjected to whole-genome sequencing alongside the inoculum. Briefly, isolates were recovered on selective agar plates grown under anaerobic conditions, and single colonies were inoculated into pre-reduced BHI-S media and grown overnight at 37 °C. Genomic DNA was extracted using the DNeasy Blood and Tissue kit (Qiagen) according to manufacturer’s instructions and submitted for whole-genome sequencing (Plasmidsaurus). Assemblies were compared to ETBF^Tet^ VPI-13784 inoculum to identify single nucleotide polymorphisms using Mauve analysis implemented Geneious Prime^103^).

For fecal Western blot analyses, fecal suspensions were centrifuged at 20,000 ×*g* for ten minutes. To detect MB03 from *B. ovatus*^Erm^–MB03 treated animals, fecal supernatant was mixed with an equal volume of 4× Laemmli loading buffer, heated at 95 °C for 10 min, and centrifuged at 20,000 × *g* for three minutes. Samples were subsequently analyzed by Western blotting using an α–VSV-G antibody (Novus Biologicals) as described above.

To detect MB03-ds-derived peptides in fecal samples from mice receiving oral minibinder treatment, clarified fecal collected during the final four weeks of the experiment were pooled and subjected to affinity enrichment. Briefly, 20 μL Ni-NTA resin (Qiagen) was pre-loaded with 1 mg mL^-1^ BFT-1–H_8_ in binding buffer (500 mM NaCl, 50 mM Tris-HCl, pH 7.5, 10 mM imidazole, 0.01% Tween-20) for 30 min with rotation at 4 °C. Pooled fecal supernatants were incubated with the resin for 30 min at 4 °C with rotation to capture BFT-binding proteins. Resin was washed five times (500 mM NaCl, 50 mM Tris-HCl, pH 7.5, 10 mM imidazole), then processed for on-bead proteomic analysis as previously described^90,104^. Proteins were digested with 10 μL of 10 ng μL^-1^ sequence-grade trypsin (Promega) for six hours at 37 °C with gentle agitation. Resin was then washed twice with 20 mM ammonium bicarbonate, with supernatants collected and combined. Supernatants containing peptides were reduced with 1 mM tris(2-caboxyethyl) phosphine hydrochloride for one hour at 37 °C, alkylated with 10 mM iodoacetamide for 30 min at room temperature in the dark, and quenched with 6 mM 1,4-dithiothreitol. Trifluoroacetic acid (TFA) was added to final concentrations of 0.5% (w/v) and stored overnight at -80 °C. The following day, peptides were purified with in-house prepared stop-and-go extraction tips (StageTips)^105^ embedded with four layers of Empore™ styrene divinyl benzene (SDB-RPS) extraction disks (Sigma). StageTips were sequentially conditioned with 100% methanol, 100% acetonitrile (ACN), 75% ACN and 5% ammonium hydroxide, 75% ACN and 0.5% acetic acid, and finally 0.1% TFA. Peptide samples were loaded onto conditioned StageTips then sequentially washed with 0.1% TFA, 75% ACN and 0.5% acetic acid, 0.5% acetic acid, and eluted with 75% ACN and 5% ammonium hydroxide. Samples were dried using a SpeeedVac^TM^ vacuum concentrator (Thermo Fisher Scientific) and resuspended in 5% ACN and 5% formic acid and analyzed by LC-MS/MS on a Lumos Fusion Orbitrap mass spectrometer (Thermo Scientific) as previously described^106^. Data were analyzed using MaxQuant^107^.

## Acknowledgements

We thank Josh Woodward for providing the negative control minibinder sequence and colon tissue for organoid generation, Maureen Thomason for assistance with organoid harvesting, Lieselotte Kreuk for support with mouse studies, Cynthia Sears, Maxwell White and Jessica Queen for providing BFT-2 and claudin-4-related reagents, Florian Hladik for providing HT-29 cells, the Yale Comparative Pathology core and Carmen Booth for pathology analyses, Ricard Rodriguez-Mias for mass spectrometry assistance, Saman Fatima for help with minibinder stability assays, Luki Goldschmidt for help with compute infrastructure, and Kaitlin Schaefer, Simon Dove, Andy Goodman, Charles Craik, Tyler Detomasi, and Mougous and Bhardwaj lab members for helpful discussions. This work was supported by the Howard Hughes Medical Institute (HHMI) Emerging Pathogens Initiative (to J.D.M, G.B., A.A.W.), the Washington Research Foundation Fellowship (P.S.), NIH R01AI170835, R01AI189789, and R01AI139087 (M.D.), the University of Washington Post-baccalaureate Research Education Program and R25GM086304 (N.H.). Crystallographic diffraction data were collected at the Advanced Photon Source (APS) at the Northeastern Collaborative Access Team beamlines funded by NIH-ORIP HEI grant S10OD021527, and the National Synchrotron Light Source II (NSLS-II), supported by the Center for BioMolecular Structure (CBMS). The APS is a U.S. Department of Energy (DOE) Office of Science User Facility operated for the DOE Office of Science by the Argonne National Laboratory under Contract No. DE-AC02-06CH11357. CBMS is supported by the National Institute of General Medical Sciences from the National Institutes of Health (P30GM133893) and by the DOE Office of Biological and Environmental Research (KP1605010). NSLS-II is a U.S. DOE Office of Science User Facility operated for the DOE Office of Science by Brookhaven National Laboratory under Contract No. DE-SC0012704. J.D.M. is an HHMI investigator, held the Lynn and Michael Garvey Chair in Gastroenterology at the University of Washington, and currently holds the John F. Enders Professorship in Microbial Pathogenesis at Yale University.

## Supplemental Figure Legends

**Supplemental Figure 1.**
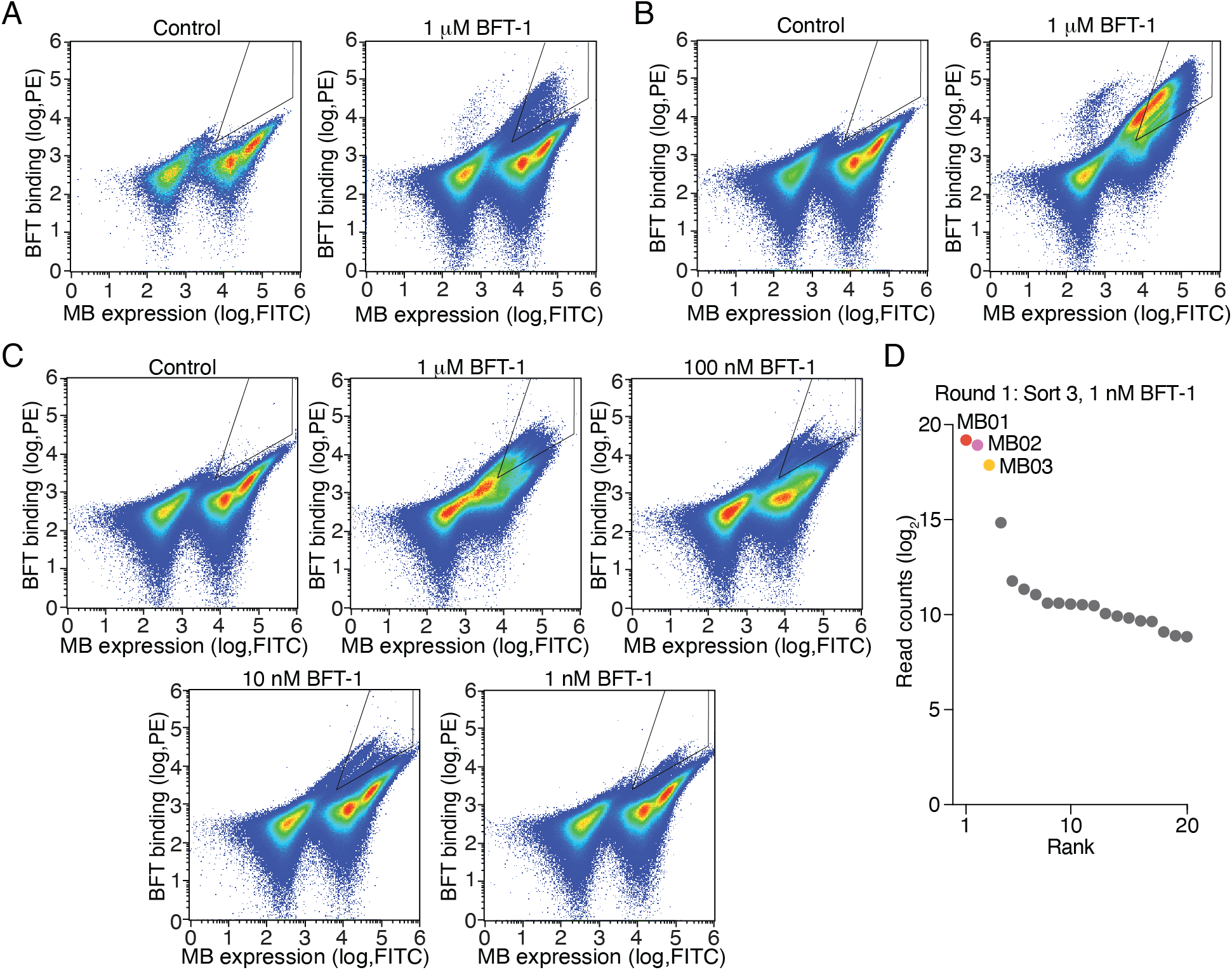
Yeast surface display-based screening of minibinders designed against the BFT-1 active site. A-C) Pseudocolor density plots of FITC and PE fluorescence from round one (A) and round two yeast display minibinder library enrichment with exposure to 1 mM BFT-1 (right) or the control treatment, and round three of enrichment (C) which employed a titration of BFT-1 from 1 nM to 1 mM. D) Relative enrichment of yeast clones encoding the indicated minibinders from the third FACS-based enrichment for BFT-1 binders (1 nM BFT-1 sample), as determined by library sequencing read counts.

**Supplemental Figure 2.**
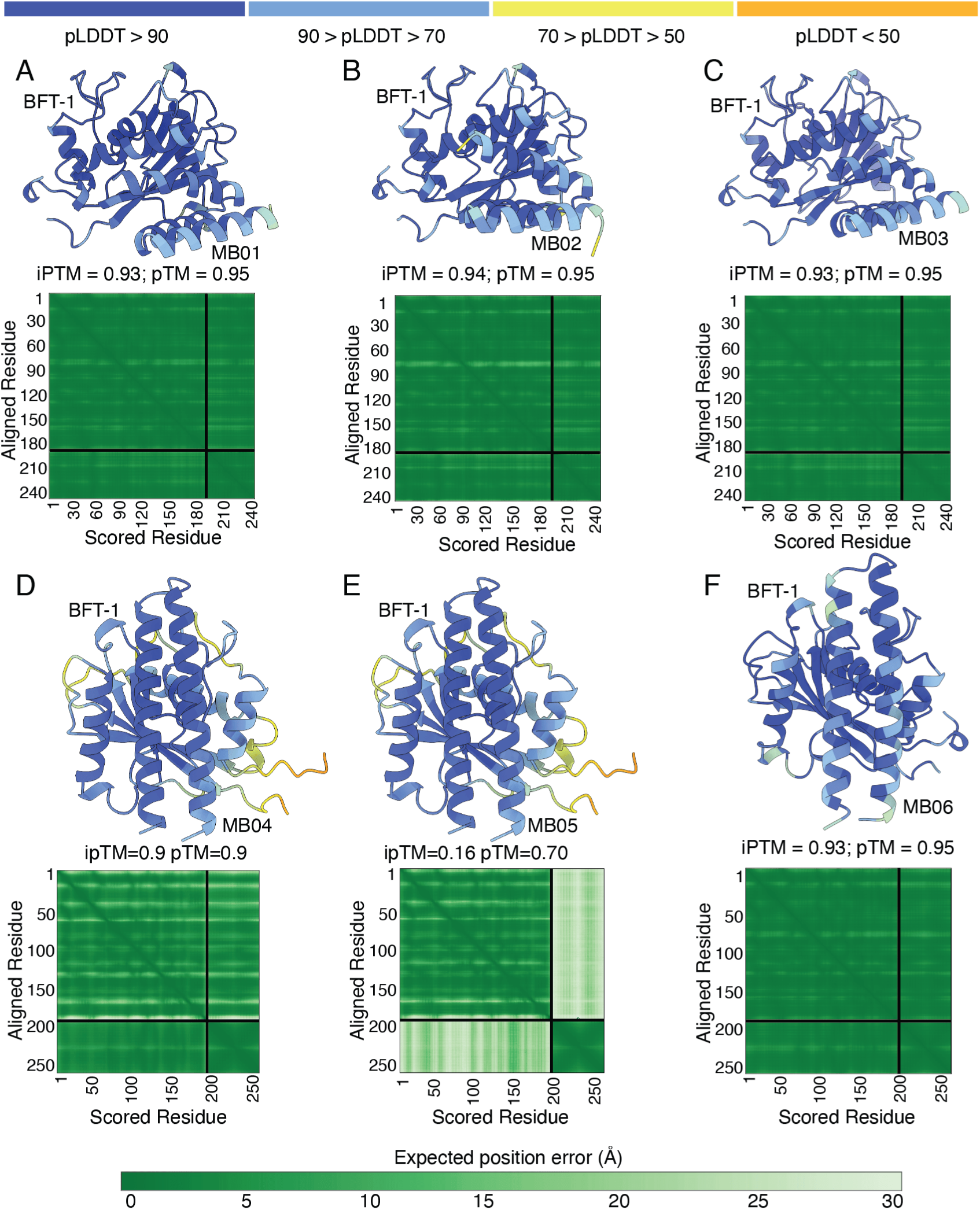
First generation BFT-1 minibinders enriched from yeast surface display are predicted to interact with two surfaces of BFT-1. A-E) AF3 models for complexes formed by BFT-1 and the indicated minibinders, colored by predicted local distance difference test (pLDDT) values. Corresponding predicted aligned error plots shown underneath each model.

**Supplemental Figure 3.**
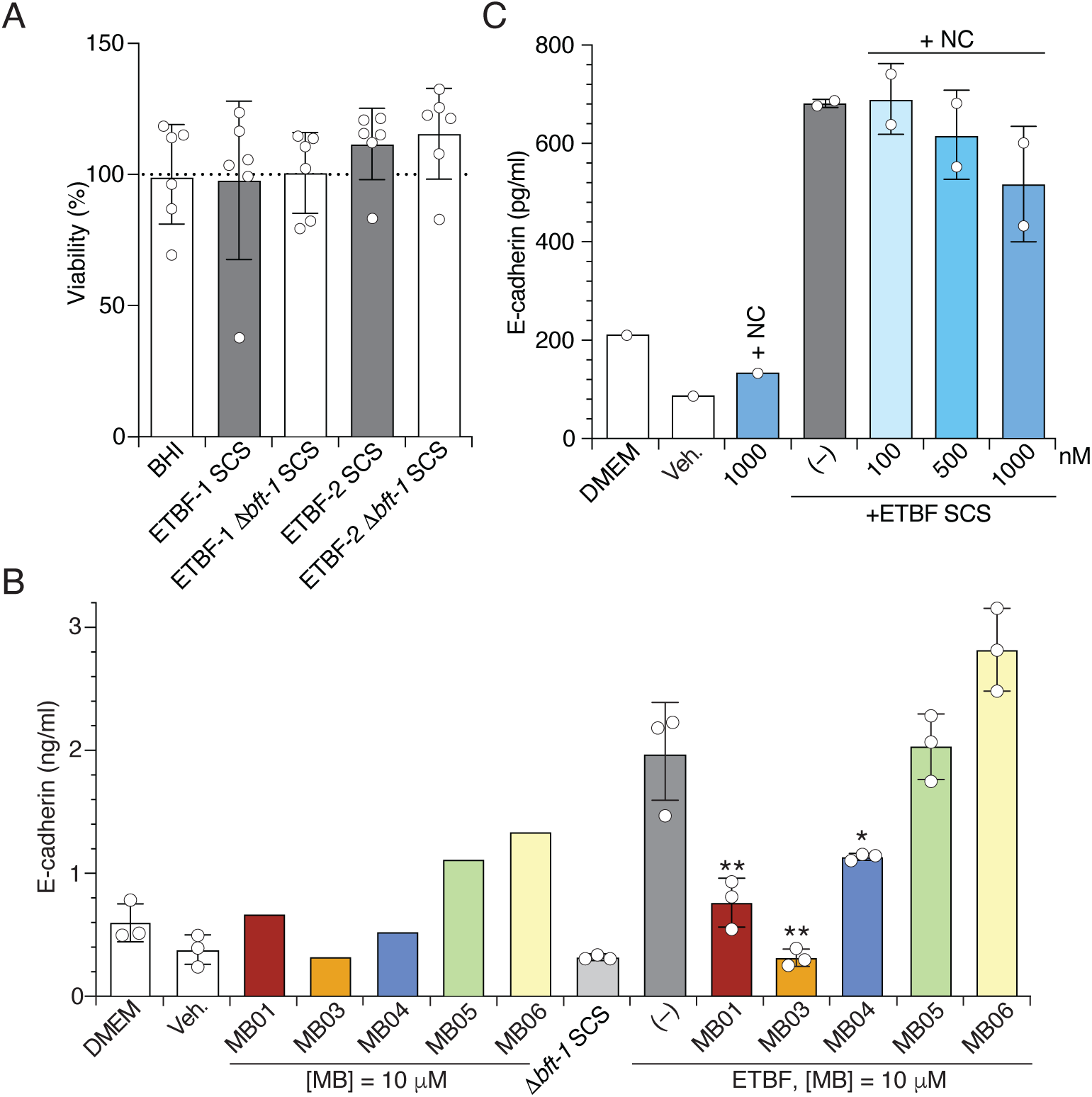
Viability of HT-29 cells is unaffected by ETBF spent culture supernatant, and the BFT inhibitory activity of PIBs is specific. A) Viability of HT-29 cells (as assessed by MTT assays) exposed SCS derived from the indicated strains, or media alone B,C) E-cadherin detected in the cell supernatant from HT-29 cells exposed to minibinders alone, ETBF SCS, or both (B), or ETBF and increasing concentrations of negative control (NC) minibinder designed against a different target (C). Data represent means and standard errors. Asterisks in B indicate minibinder treatments significantly different from ETBF SCS treatment alone (1-way ANOVA with Dunnett’s multiple comparisons test, *p<0.01, **p<0.001, ***<0.0001).

**Supplemental Figure 4.**
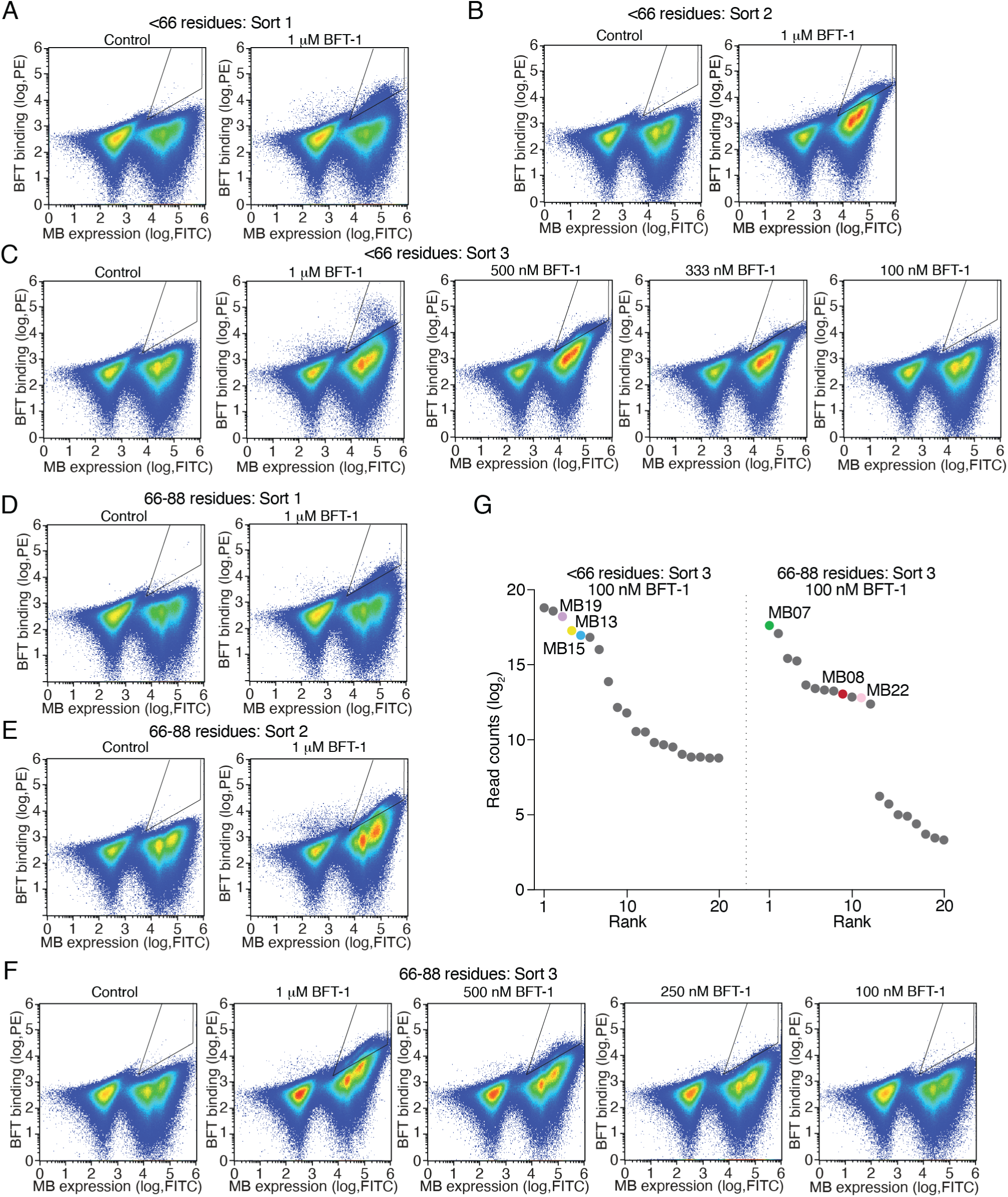
Yeast surface display-based screening of minibinders designed against the prodomain interface of BFT-1. A-F) Pseudocolor density plots of FITC and PE fluorescence from <66 residue design library (A-C) and the 66-88 residue design library (D-F) after one (A,D) or two (B,E) rounds of exposure to 1 mM BFT-1 (right) or the control treatment (left) or third enrichment employing a titration of BFT concentrations (C,F). G) Relative enrichment of yeast clones encoding the indicated minibinders from the third FACS-based enrichment for BFT-1 binders (100 nM BFT-1 sample), as determined by library sequencing read counts.

**Supplemental Figure 5.**
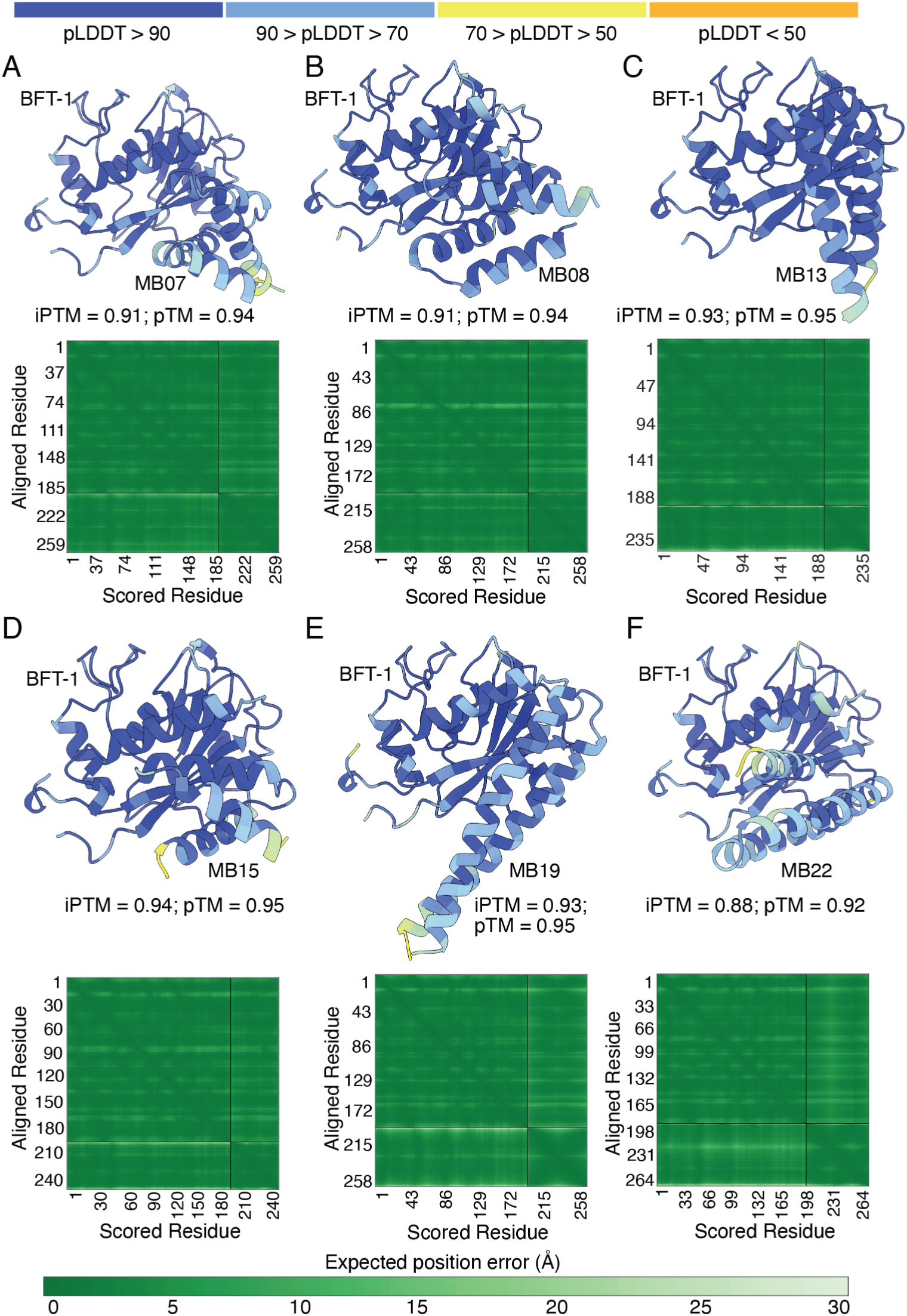
Second generation minibinders are predicted to interact with the prodomain of BFT-1. A-E) AF3 models for complexes formed by BFT-1 and the indicated minibinders, colored by predicted local distance difference test (pLDDT) values. Corresponding predicted aligned error plots shown underneath each model.

**Supplemental Figure 6.**
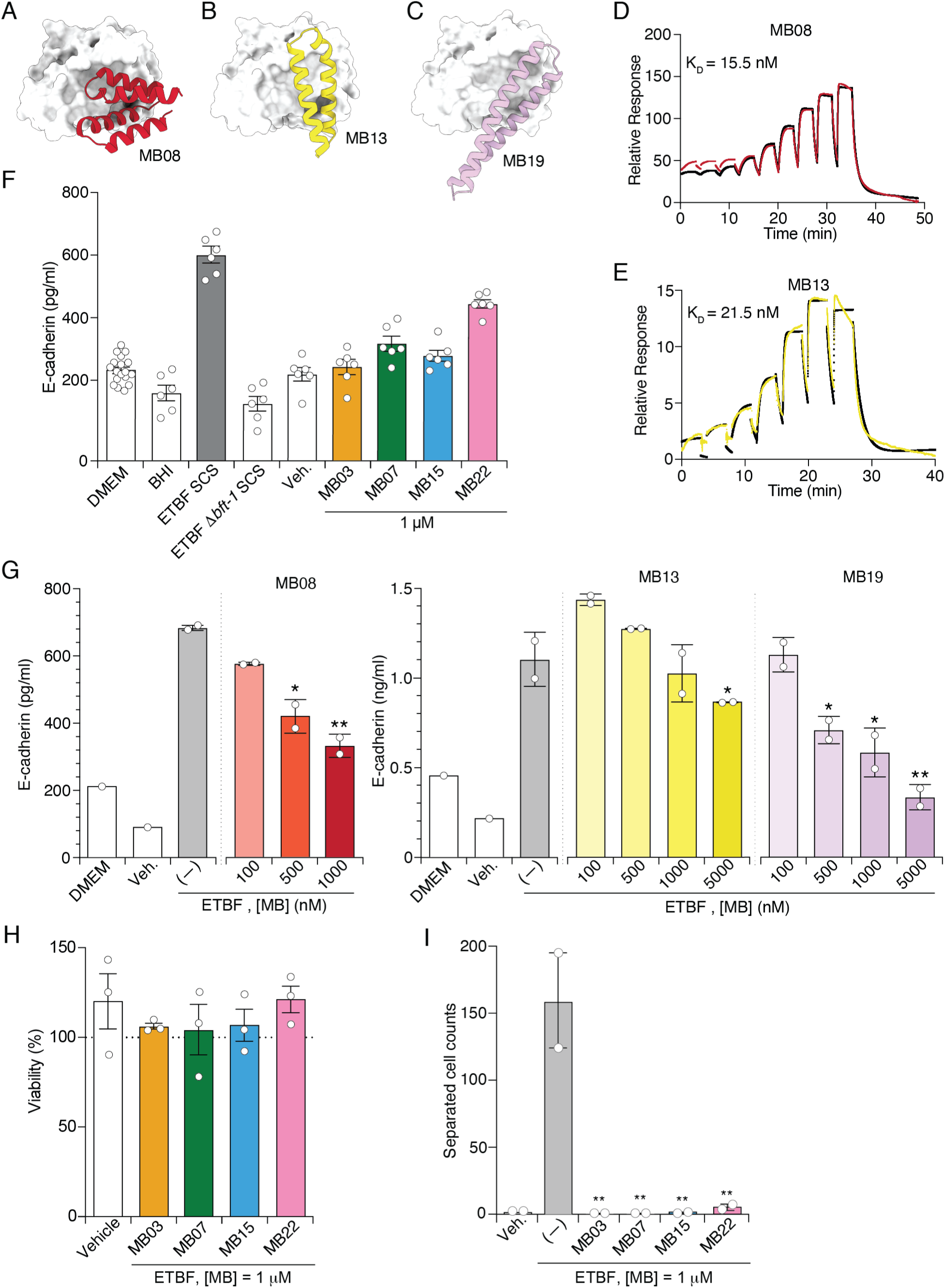
Second generation BFT inhibitors perform comparably to PIB MB03. A-C) AF3 models of the BFT-1 (white, surface display) interaction with second generation minibinders that passed initial screening criteria but performed poorly in one or more experimental assay. D, E) SPR analysis of the binding affinity of the indicated minibinders for immobilized BFT-1. K_D_ values indicate binding affinity calculated from this analysis. F,G) E-cadherin detected in the cell supernatant from HT-29 cells exposed to the indicated purified minibinders alone (F), with ETBF SCS (G) or the indicated controls. H) HT-29 viability (MTT) assays of cells exposed to the indicated purified minibinders. I) Counts of separated, Giemsa-stained HT-29 cells exposed to the indicated minibinders and ETBF SCS or a vehicle control.

**Supplemental Figure 7.**
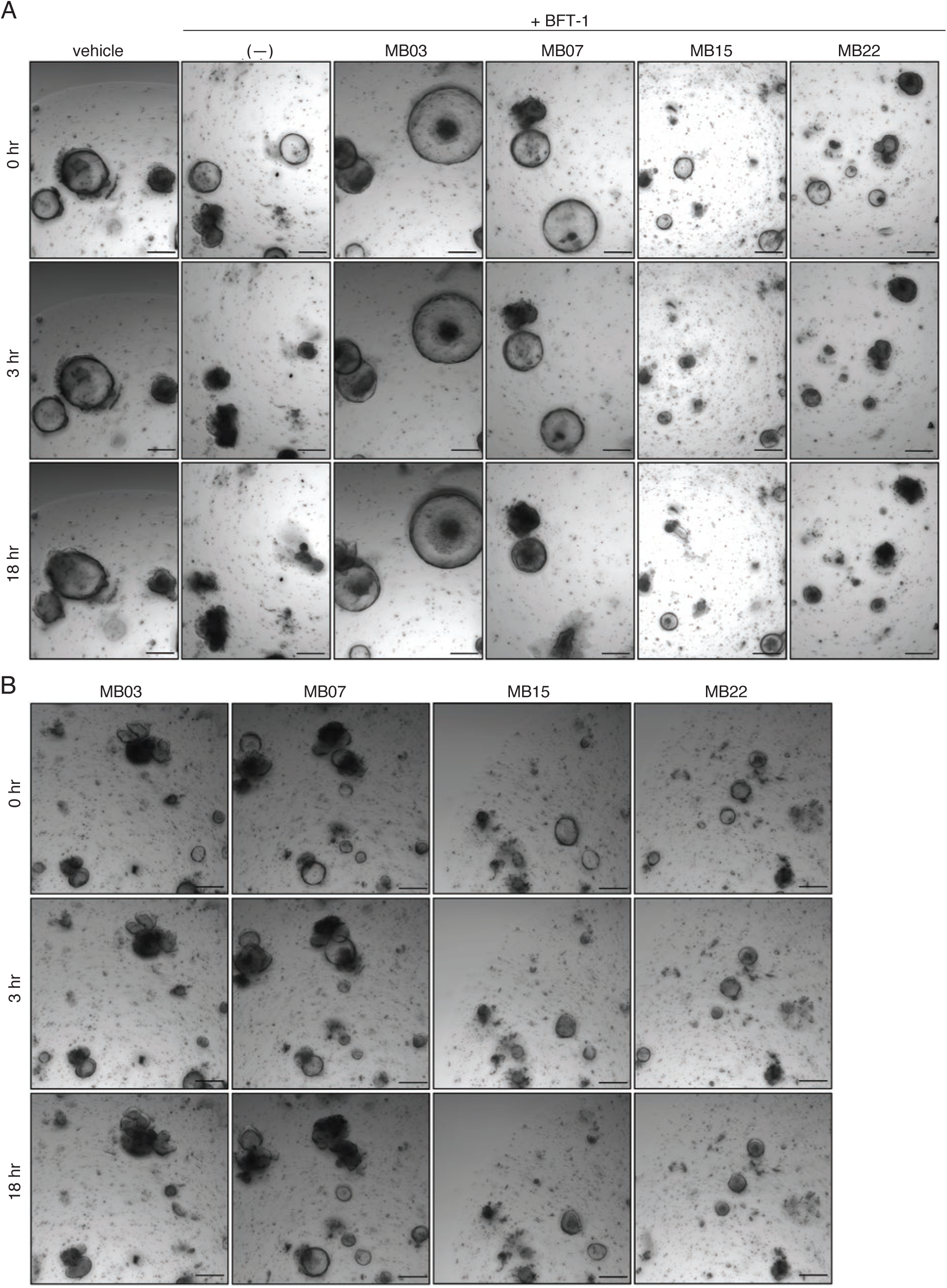
PIBs prevent BFT-1-mediated murine colonoid damage. A, B) Murine colonoid integrity following treatment with purified BFT-1 (100 nM) with the indicated minibinders (10 mM) (A), or the indicated minibinders alone (B).

**Supplemental Figure 8.**
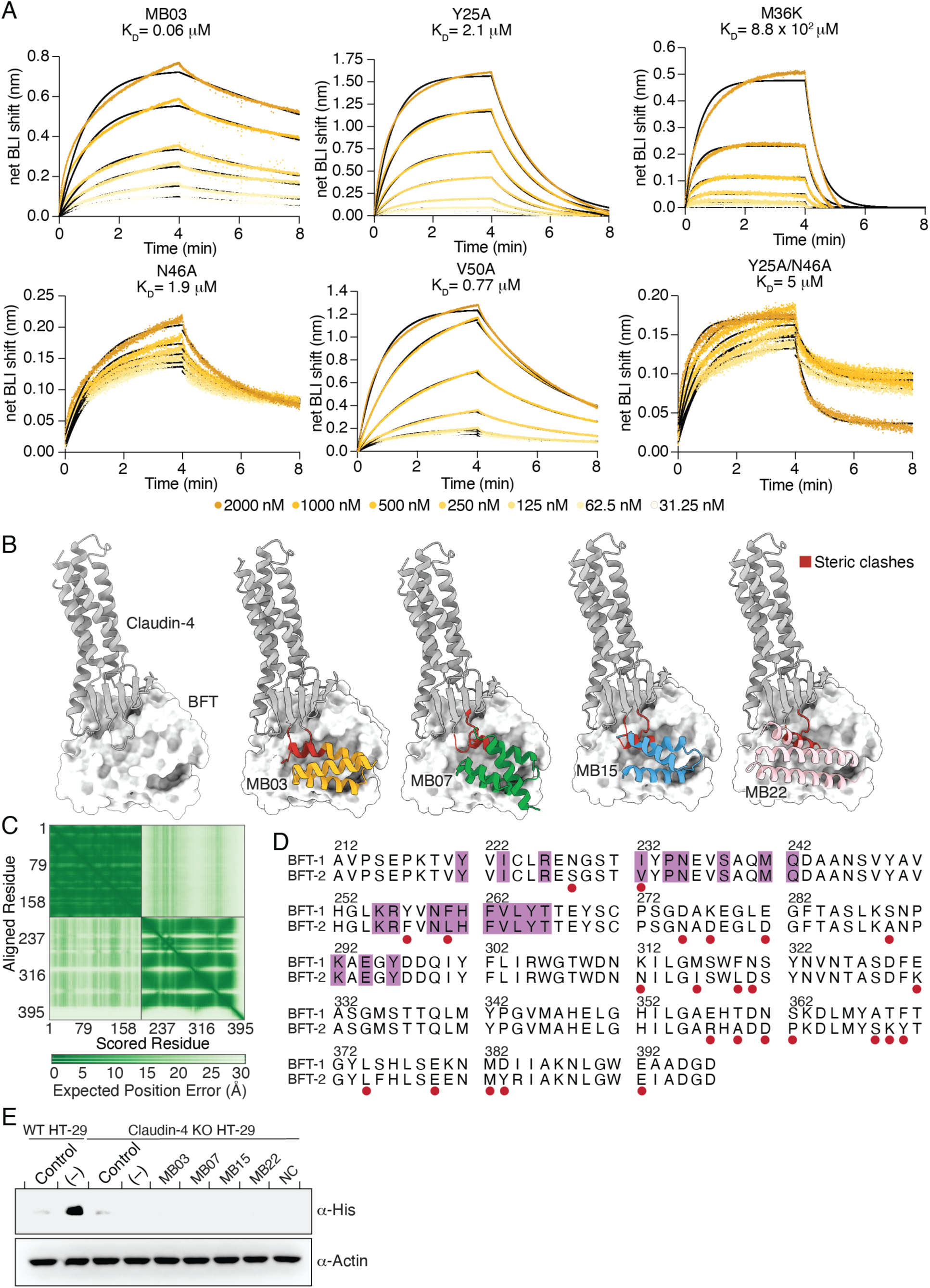
Binding assays support predicted MB03-BFT-1 interaction residues, and PIBS interfere with the BFT-claudin-4 interaction. A) Biolayer interferometry analysis of the binding affinity of BFT-1 for immobilized MB03 variants containing the indicated substitutions. K_D_ values indicate binding affinity calculated from this analysis. Colored traces indicate measured data; black curves indicate corresponding fitted data. B) AF3 model of BFT-1 (surface) to claudin-4 (ribbon, grey), with and without the indicated minibinder. Steric clashes are defined using the MolProbity criterion (interatomic distance is less than the sum of their van der Waals radii minus 0.4 Å) and are shown in red. C) Predicted aligned error plot of BFT-claudin-4 interaction model. D) Sequence alignment of the BFT-1 and BFT-2 catalytic domains. Prodomain interface (purple) and non-conserved (red) residues indicated. E) Western blot analysis of proteins extracted from WT or claudin-4 KO HT-29 cells alone (control) or incubated with BFT-2–H_6_ (5 nM) and the indicated PIB (500 nM).

**Supplemental Figure 9.**
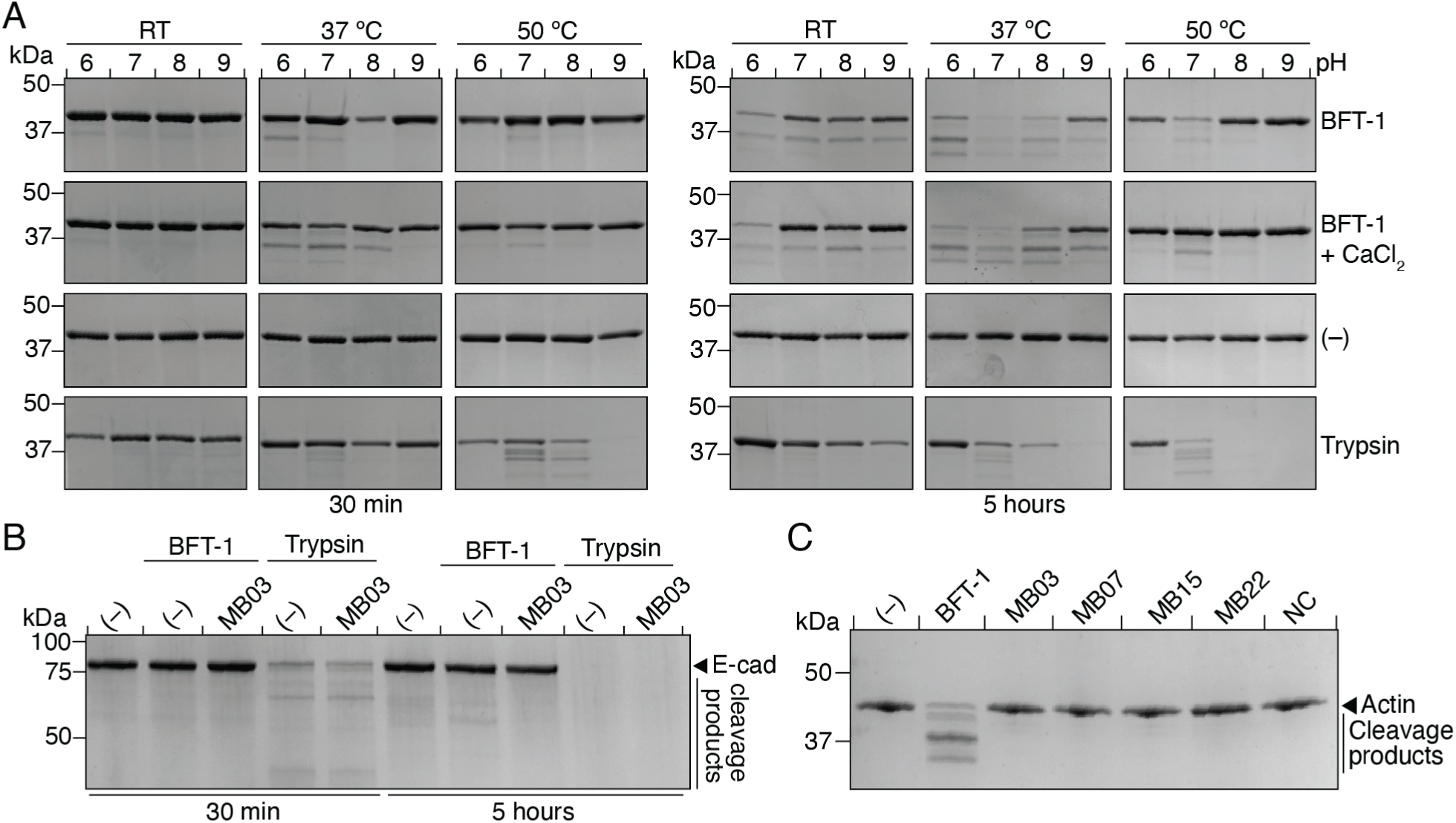
Optimization of assays conditions to detect catalytic activity of BFT-1. A-C) Coomassie-stained SDS-PAGE analysis of cleavage products generated from actin (2 mM, A, C) or E-cadherin (2 mM, B) incubated with purified BFT-1–H_8_ (20 nM), no enzyme (–, negative control), trypsin (20 nM, positive control), or the indicated purified minibinders alone under the noted conditions (CaCl_2_ supplied at 1 mM where indicated). Minibinders, when added, were supplied at 1 µM.

**Supplemental Figure 10.**
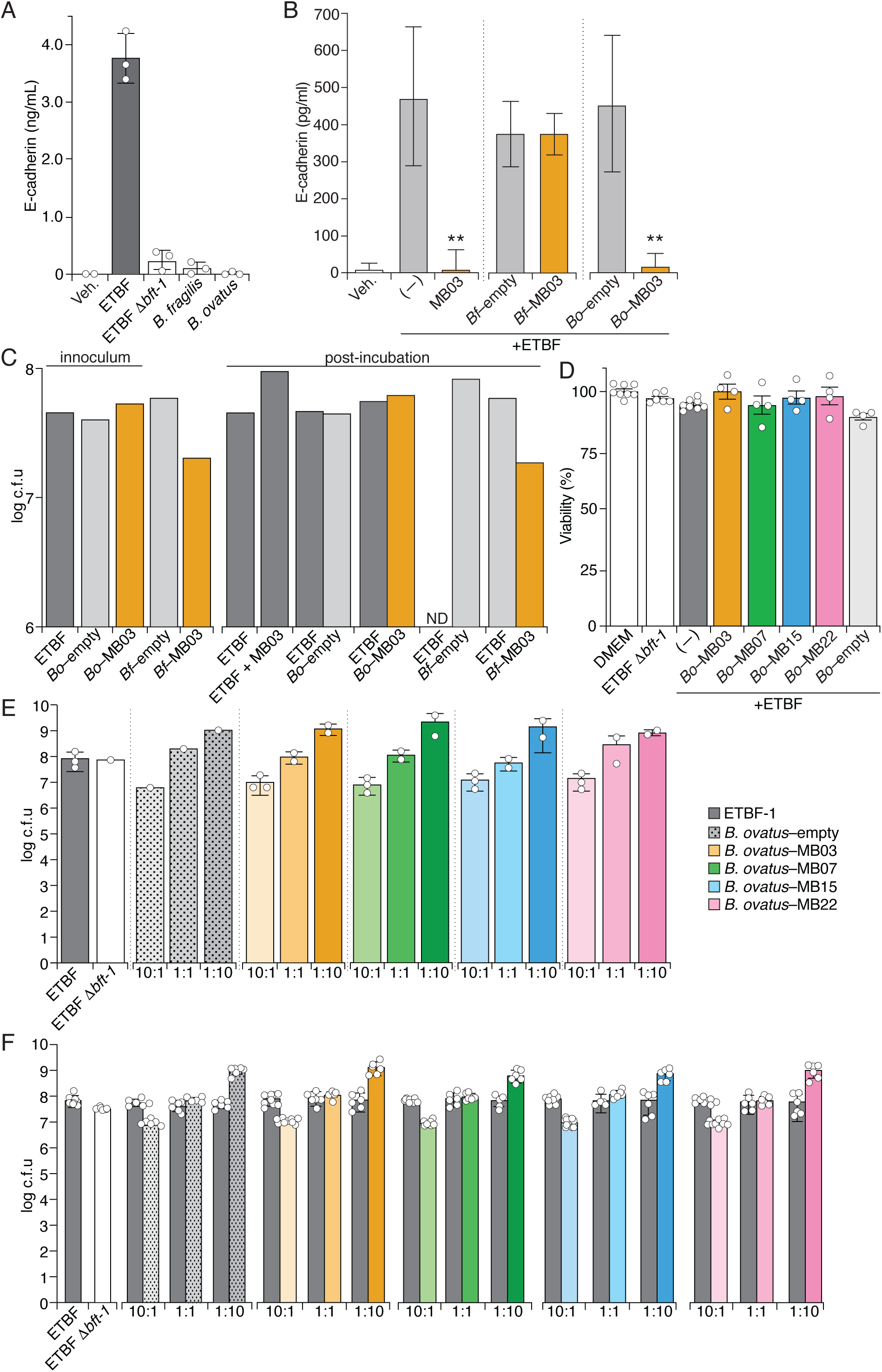
PIB-secreting strains of *B. ovatus* do not affect ETBF or HT-29 viability in co-culture assays. A,B) Released E-cadherin detected in the cell supernatant from HT-29 cells exposed to the indicated live bacteria for five hours (A) or bacterial co-cultures added at a 1:1 ratio (B). MB03 (1 µM) treatment was included for comparison. C) C.f.u counts of bacteria from one of the replicate samples of the experiment shown in (B). Dark grey, ETBF; light grey *B. ovatus*–empty or *B. fragilis*–empty; orange, MB03-secreting strains. Adjacent bars indicate populations measured in the same mouse. ETBF populations could not be assessed in the *B. fragilis-*empty colonized mouse due to a technical error. D) Viability (MTT) assays of HT-29 cells treated with the indicated live bacteria. E, F) C.f.u. counts of the bacterial inoculum (E) or post incubation with HT-29 (F). Bacteria were administered at the indicated initial ratios (ETBF:*B. ovatus*). Counts correspond to cultures assayed for E-cadherin release in Fig. 4D.

**Supplemental Figure 11.**
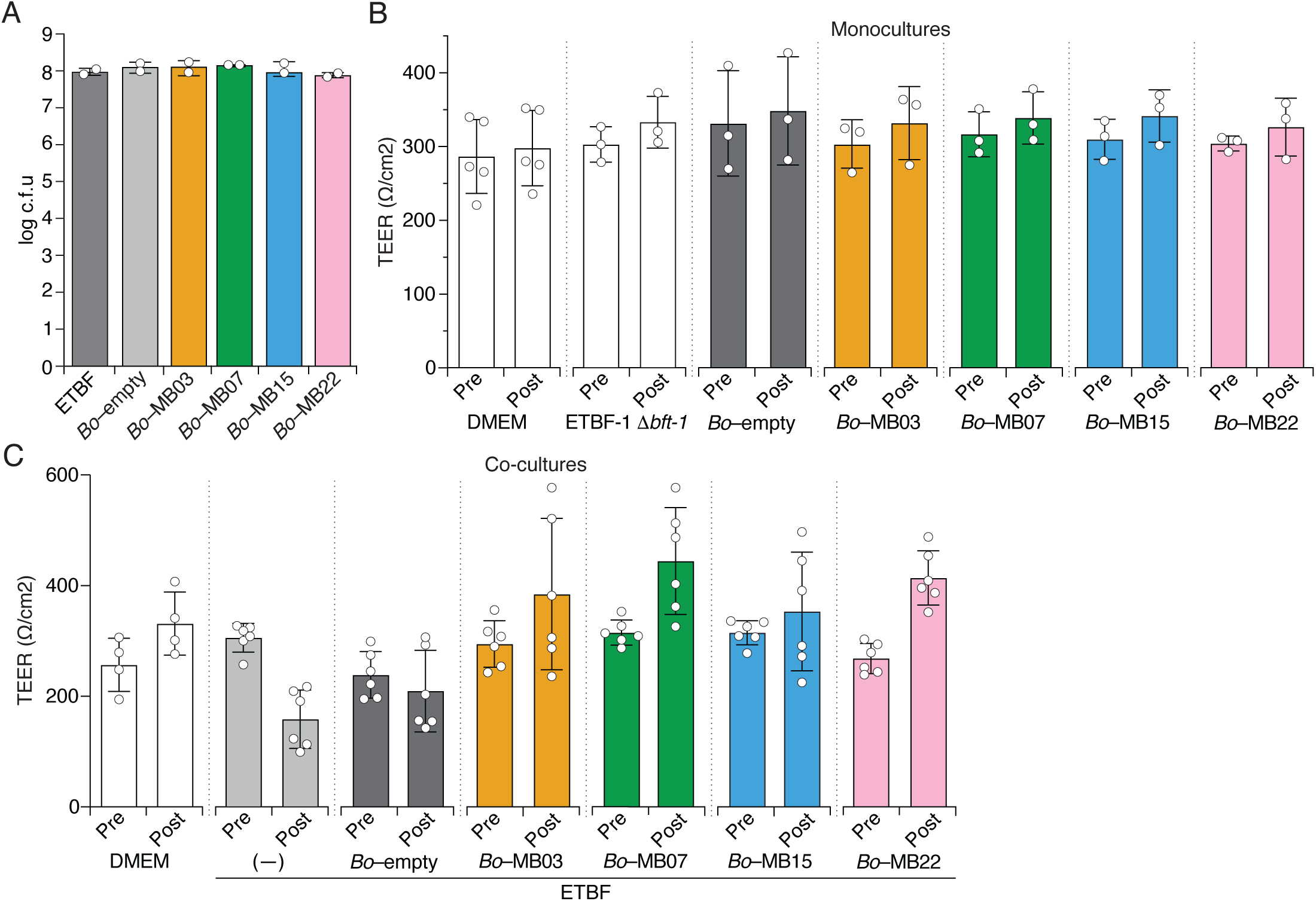
PIB-secreting *B. ovatus* strains prevent ETBF-mediated TER reductions when applied to confluent cultured epithelial cells. A) C.f.u counts of bacteria applied to the apical surface of HT-29-MTX-E12 cells grown in transwells for conducting TEER assays. B,C) TEER readings of HT-29-MTX-E12 cells treated with either bacterial monocultures (B) or co-culture (C). Co-cultures were added at a 1:1 ratio and incubated for five hours between measurements.

**Supplemental Figure 12.**
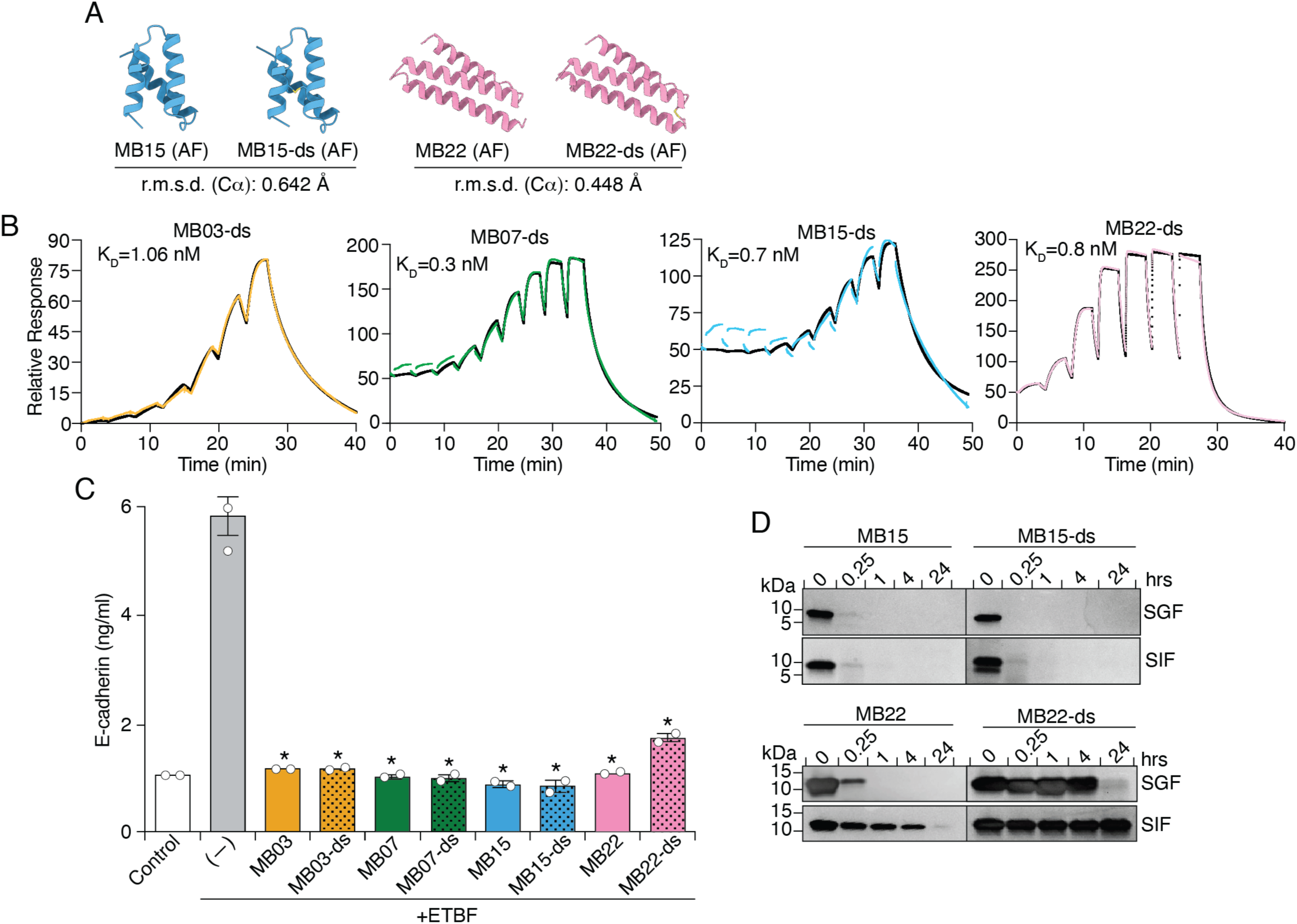
Introduction of a stabilizing disulfide bond does not substantially reduce PIB binding affinity or inhibition of BFT-1. A) AF3 models of the indicated minibinders. B) SPR analysis of the binding affinity of the indicated minibinders for immobilized BFT-1–H_8_. K_D_ values indicate binding affinity calculated from this analysis. C) Released E-cadherin detected in the cell supernatant from HT-29 cells exposed to the indicated purified minibinders. D) SDS-PAGE analysis of the indicated PIBs incubated for the indicated intervals in synthetic gastric or intestinal fluid (SGF, SIF). Asterisks in C indicate minibinder treatments significantly different from ETBF SCS treatment alone (1-way ANOVA with Dunnett’s multiple comparisons test, *p<0.0001). Data represent means and standard errors. Asterisks in C indicate minibinder treatments significantly different from ETBF SCS treatment along (1-way ANOVA with Dunnett’s multiple comparisons test, * *<0.0001).

**Supplemental Figure 13.**
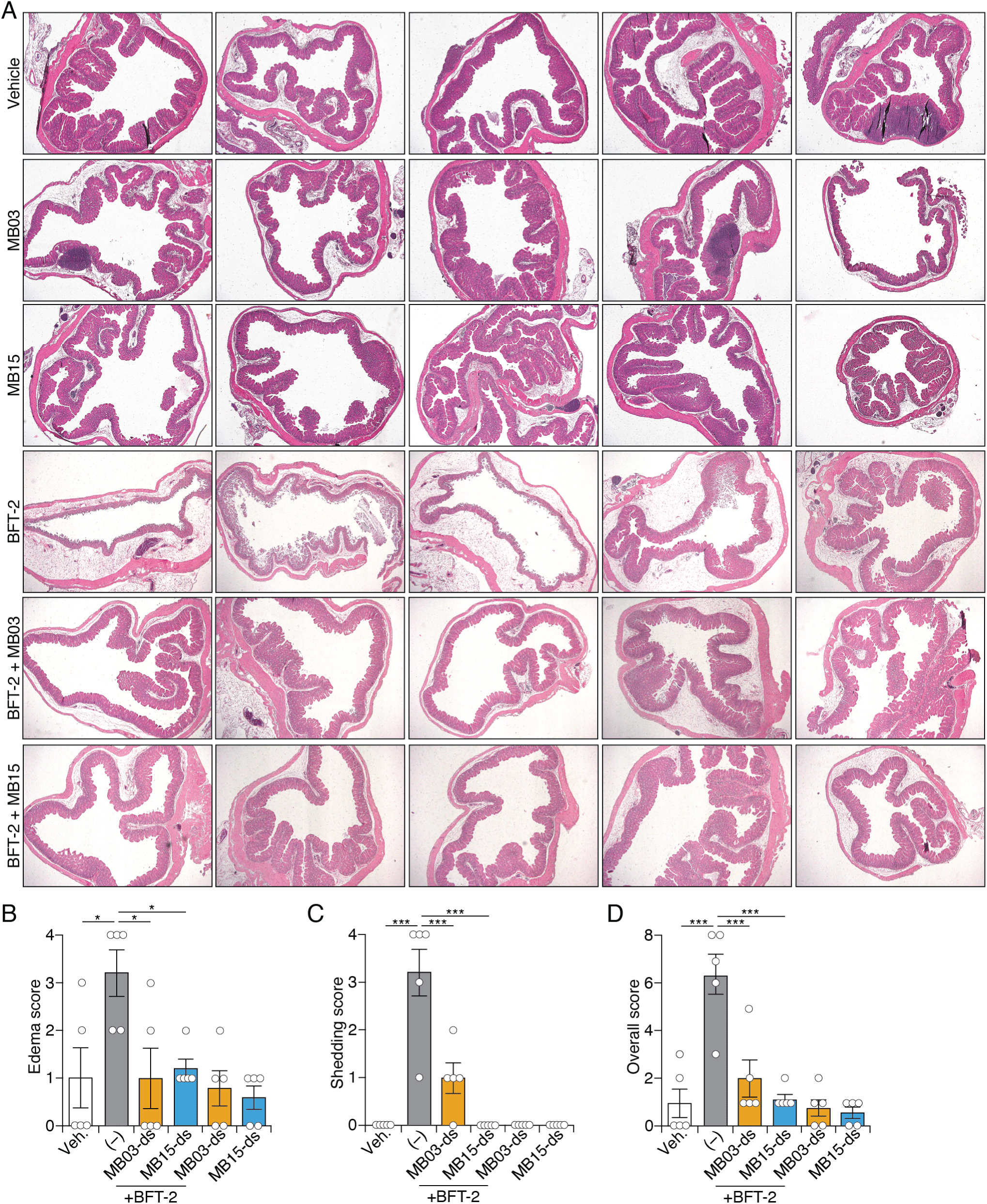
Mouse cecal injection data. A) Full hematoxylin and eosin (H&E)-stained sections of mouse ceca injected with either PBS (vehicle), MB03-ds, MB15-ds, BFT-2, BFT-2 and MB03-ds, or BFT-2 and MB15-ds. Quantification of cecum histopathology of edema score (B), shedding score (C), and overall score (D), from ceca in (A). (1-way ANOVA with Dunnett’s multiple comparisons test, *p<0.05, ***<0.001).

**Supplemental Figure 14.**
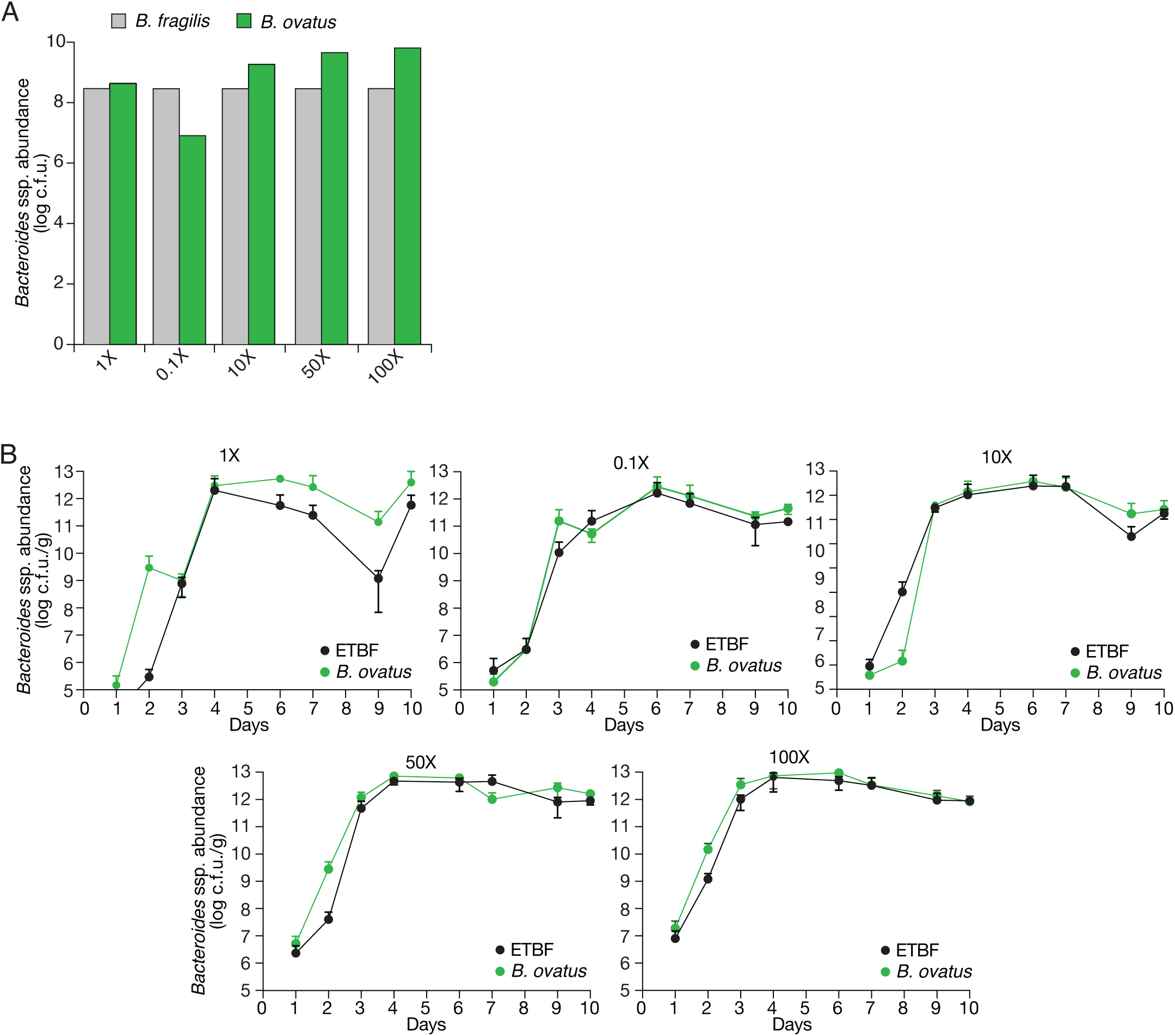
Mice are successfully co-colonized with ETBF and *B. ovatus*. A) C.f.u. counts of bacterial inocula administered to antibiotic pre-treated mice by oral gavage during colonization experiments shown in B. B) Fecal abundance of the indicated strains from mice colonized with ETBF monoculture and *B. ovatus* administered at the indicated ratio of *B. ovatus* to ETBF (0.1-100X). Strains contained antibiotic resistance cassettes (erm and tet, respectively) for quantification by plating on selective media. Graphs show the mean and standard deviation of three to six mice per group.

**Supplemental Figure 15.**
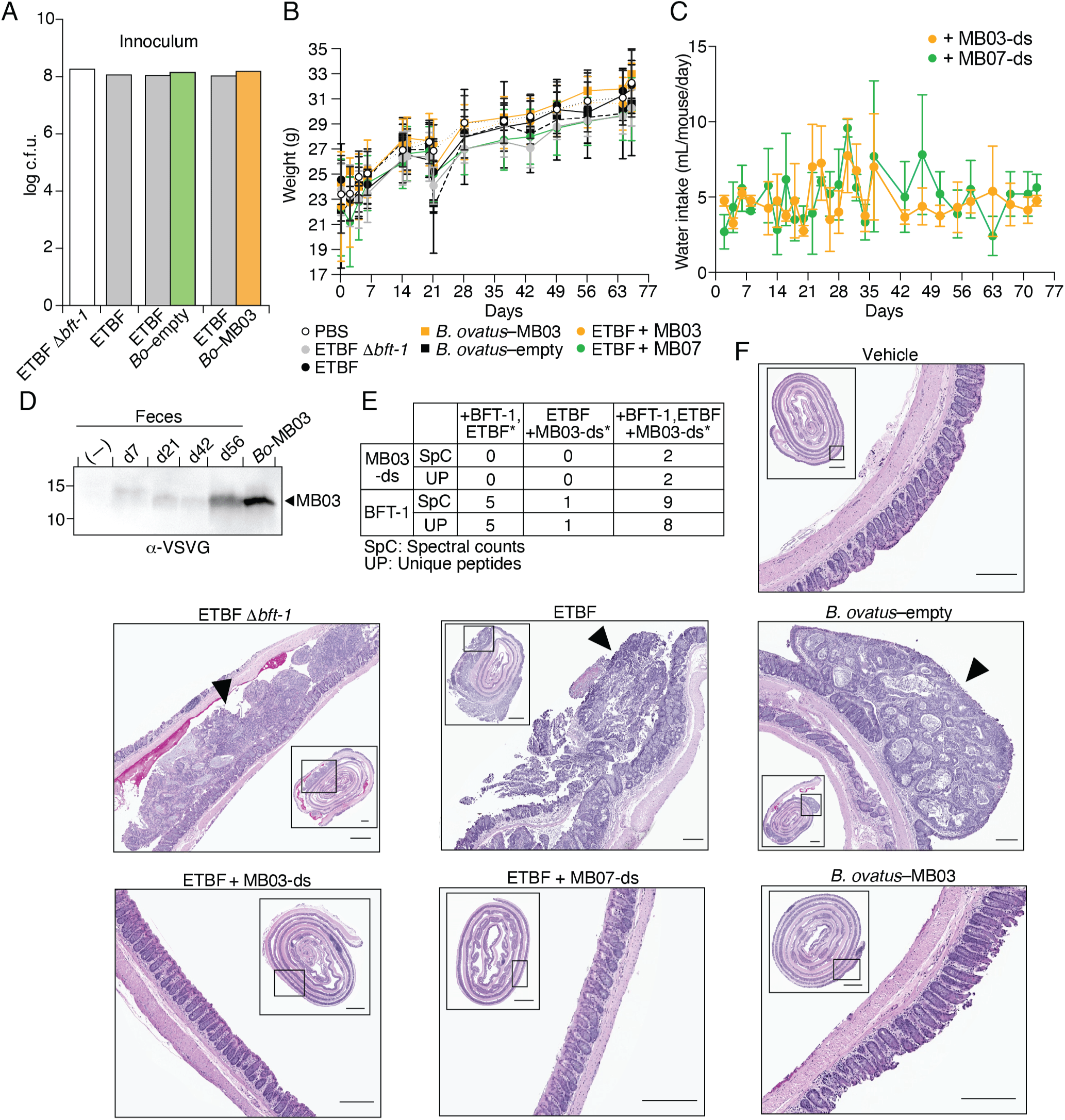
AOM/DSS treated mice with LBPs or PIBs in drinking water have reduced tumor burden. A) C.f.u. counts of bacterial inoculum administered to mice by oral gavage. B) Mean body weights of mice exposed to the indicated treatments. C) Water consumption of the minibinder-treated mice throughout the experiment. Plots in B and C indicate means and standard errors derived from three to ten mice per group. D) Western blot analysis of VSV-G–MB03 in fecal pellets collected from representative mice the indicated number of days post introduction of *B. ovatus–*MB03 via oral gavage. E) Spectral counts and unique peptides detected by mass spectrometry analysis of proteins associated with Ni-NTA beads incubated with BFT–H_8_ (or beads only control, middle) and clarified extracts of pooled fecal pellets collected from mice colonized with ETBF (first column) or colonized with ETBF and supplied with MB03-ds in drinking water (last column). F) Representative hematoxylin and eosin (H&E)-stained colons from the indicated treatment groups. Insets show intact Swiss rolled colons, with the enlarged regions, representing distal colons, boxed. Black arrows indicate tumors. Inset scale bar = 1 mm; enlargement scale bar = 200 μm.

