## Supplemental Table 1 for "Orally available designed miniproteins inhibit enterotoxigenic *Bacteroides fragilis* pathology by blocking toxin receptor binding"

Supplementary Table 1. Kinetic characterization of BFT minibinders

| **Minibinder** | **K_D_ (nM)** | **K_off_ (s^-1^)** |
| --- | --- | --- |
| G1-MB01 | 20 | 6.14E-02 |
| G1-MB03 | 0.66 | 1.66E-03 |
| G2-MB07 | 11.5 | 7.80E-03 |
| G2-MB08 | 15.5 | 3.39E-02 |
| G2-MB13 | 21.5 | 1.40E-02 |
| G2-MB15 | 1.45 | 1.23E-02 |
| G2-MB22 | 3.46 | 9.08E-03 |
