## Supplemental Table 2 for "Orally available designed miniproteins inhibit enterotoxigenic *Bacteroides fragilis* pathology by blocking toxin receptor binding"

Supplementary Table 2. Data collection and refinement statistics for minibinder crystal structures

|  | **MB03-ds**  **(PDB ID: 36LX)** | **MB07**  **(PDB ID: 36LY)** | **MB07-ds**  **(PDB ID: 36MB)** |
| --- | --- | --- | --- |
| Space Group | P 21 21 21 | P 1 | P 1 21 1 |
| Resolution (Å) | 40  - 2.1 (2.18  - 2.1)* | 32.97  - 2.55 (2.92  - 2.55) | 56  - 2.0 (2.05  - 2.0) |
| Cell dimensions |  |  |  |
| a, b, c (Å) | 44.265, 57.412, 93.432 | 33.042, 34.711, 37.038 | 55.306 51.306 56.008 |
| α, β, γ (º) | 90, 90, 90 | 81.61, 70.99, 72 | 90 90.95 90 |
| Total reflections | 131257 (7707) | 18588 (6163) | 179854 (11738) |
| Unique reflections | 17587 (1498) | 6771 (2241) | 26405 (1891) |
| Multiplicity | 7.5 (5.1) | 2.7 (2.8) | 6.8 (6.2) |
| R merge | 0.1488 (1.897) | 0.1497 (0.9153) | 0.24 (1.361) |
| R pim | 0.05248 (0.8244) | 0.1128 (0.6943) | 0.09753 (0.5856) |
| CC1/2 | 0.994 (0.456) | 0.966 (0.618) | 0.985 (0.566) |
| CC* | 0.999 (0.791) | 0.991 (0.874) | 0.996 (0.85) |
| I/σI | 6.03 (0.71) | 2.89 (0.97) | 5.23 (0.91) |
| Completeness (%) | 97.23 (90.16) | 94.58 (94.05) | 99.10 (99.41) |
| *Refinement* |  |  |  |
| Reflections used in refinement | 14077 (1273) | 4558 (1518) | 21207 (1622) |
| Reflections used for R-free | 1408 (127) | 455 (147) | 2010 (155) |
| R-work | 0.2534 (0.3122) | 0.2225 (0.2573) | 0.2267 (0.2919) |
| R-free | 0.2886 (0.3560) | 0.2830 (0.3440) | 0.2663 (0.3477) |
| *Ramachandran Statistics* |  |  |  |
| Favored (%) | 99.11 | 95.95 | 100 |
| Allowed (%) | 0.89 | 4.05 | 0 |
| Outliers (%) | 0 | 0 | 0 |
| RMS (bonds) | 0.003 | 0.003 | 0.003 |
| RMS (angles) | 0.48 | 0.52 | 0.54 |
| Average B-factor | 36.22 | 46.43 | 38.06 |
| macromolecules | 35 | 48 | 38 |
| solvent | 34 | 36 | 42 |
| ligands | n/a | 70 | 51 |

*Values in parentheses reflect the highest-resolution shell
