## Supplemental Table 3 for "Orally available designed miniproteins inhibit enterotoxigenic *Bacteroides fragilis* pathology by blocking toxin receptor binding"

Supplementary Table 3. Intestinal tumor burden and crypt dysplasia in mice across treatment groups

| **Mouse number** | **Treatment** | **HE* Tumor Number^1,2^** | **Crypt Dysplasia^3^** |
| --- | --- | --- | --- |
| 1 | PBS | 0.00 | 0 |
| 2 | PBS | 0.00 | 0 |
| 3 | PBS | 0.00 | 0 |
| 4 | PBS | 0.00 | 1 |
| 5 | PBS | 0.00 | 0 |
| Average |  | 0.00 | 0.20 |
| Total |  | 0.00 |  |
| 1 | ETBF ∆*bft-1* | 0.00 | 0 |
| 2 | ETBF ∆*bft* | 0.00 | 1 |
| 3 | ETBF ∆*bft* | 1.00 | 1 |
| 4 | ETBF ∆*bft* | 3.50 | 1 |
| 5 | ETBF ∆*bft* | 4.00 | 1 |
| Average |  | 1.70 | 0.80 |
| Total |  | 8.50 |  |
| 1 | ETBF | 1.00 | 1.5 |
| 2 | ETBF | 1.00 | 5 |
| 3 | ETBF | 3.00 | 5 |
| 4 | ETBF | 3.50 | 5 |
| 5 | ETBF | 7.00 | 5 |
| Average |  | 3.10 | 4.30 |
| Total |  | 15.50 |  |
| 1 | *B. ovatus*-erm | 0.00 | 1 |
| 2 | *B. ovatus*-erm | 3.00 | 1 |
| 3 | *B. ovatus*-erm | 4.50 | 1 |
| Average |  | 2.50 | 1.00 |
| Total |  | 6.50 |  |
| 1 | MB03 | 0.00 | 1 |
| 2 | MB03 | 0.00 | 0 |
| 3 | MB03 | 0.00 | 0 |
| 4 | MB03 | 0.00 | 1 |
| 5 | MB03 | 0.00 | 0 |
| Average |  | 0.00 | 0.40 |
| Total |  | 0.00 |  |
| 1 | MB07 | 0.00 | 0 |
| 2 | MB07 | 0.00 | 0 |
| 3 | MB07 | 0.00 | 0 |
| 4 | MB07 | 0.00 | 1.5 |
| 5 | MB07 | 0.00 | 0 |
| Average |  | 0.00 |  |
| Total |  | 0.00 |  |
| 1 | *B. ovatus*-MB03 | 0.00 | 0 |
| 2 | *B. ovatus*-MB03 | 0.00 | 0 |
| 3 | *B. ovatus*-MB03 | 0.00 | 0 |
| 4 | *B. ovatus*-MB03 | 0.00 | 1 |
| 5 | *B. ovatus*-MB03 | 0.00 | 0 |
| 6 | *B. ovatus*-MB03 | 0.00 | 1 |
| Average |  | 0.00 | 0.33 |
| Total |  | 0.00 | 2 |

^1^Analysis was conducted by a board-certified pathologist blinded to the treatment groups.

^2^Tumor number per mouse was averaged across the number of Swiss-rolled section per slide; half-counts are also used for ambiguous lesions

^3^Crypt dysplasia scoring: present (1 = yes), absent (0 = no)
